# Cullin1 represses systematic inflammasome activation by binding and catalyzing NLRP3 ubiquitination

**DOI:** 10.1101/289637

**Authors:** Pin Wan, Qi Zhang, Weiyong Liu, Yaling Jia, Tianci Wang, Wenbiao Wang, Pan Pan, Ge Yang, Qi Xiang, Siyu Huang, Qingyu Yang, Wei Zhang, Fang Liu, Kailang Wu, Yingle Liu, Jianguo Wu

## Abstract

Activation of the NLRP3 inflammasome is a key process of host immune response, the first line of defense against cellular stresses and pathogen infections. However, excessive inflammasome activation damages the hosts, and thus it must be precisely controlled. The mechanism underlying the repression of systematic inflammasome activation remains largely unknown. This study reveals that CUL1, a key component of the SCF E3 ligase, plays a critical role in regulation of the inflammasome. CUL1 suppresses the inflammasome activation in HEK293T cells, inhibits endogenous NLRP3 in macrophages, and represses inflammatory responses in C57BL/6 mice. Under normal physiological conditions, CUL1 interacts with NLRP3 to disrupt the inflammasome assembly, and catalyzes NLRP3 ubiquitination to repress the inflammasome activation. In response to inflammatory stimuli, CUL1 disassociates from NLRP3 to release the repression of NLRP3 inflammasome activation. This work reveals a distinct mechanism underlying the repression of inflammasome activation under physiological conditions and the induction of inflammasome activation in response to inflammatory stimuli, and thus provides insights into the prevention and treatment of infectious and inflammatory diseases.

## Introduction

Innate immune response is the first line of defense against cellular stresses and pathogen infections [1]. A key process of host immunity is the activation of inflammasomes [2,3]. The NLRP3 inflammasome, a best characterized inflammasome, is crucial for acute and chronic inflammatory responses. This inflammasome consists of three components, intracellular sensor protein (NLRP3), adaptor protein (ASC), and effecter protein (Casp-1), and regulates maturation and secretion of pro-inflammatory cytokines IL-1β and IL-18, which initiate multiple signaling pathways and drive inflammatory responses [4–7]. NLRP3 contains three domains: PYRIN (PYD), NACHT (NBD), and leucine-rich repeat (LRR) [8]. ASC comprises N-terminal PYD and C-terminal CARD [9,10]. Casp-1 harbors N-terminal caspase activation and CARD, internal large domain (p20), and C-terminal domain (p10) [11–15]. During the inflammasome activation, interaction between ASC PYD and NLRP3 PYD is necessary for ASC oligomer assembly that provides a platform for pro-Casp-1 activation [16–19]. Inflammasome activation plays a key role in host immunity, but excessive and uncontrolled activation damages host by causing infectious, inflammatory, and immune diseases [20–25], and thereby it must be tightly controlled. Although negative regulations are employed by host and pathogen to inhibit the NLRP3 inflammasome [26–28], the mechanisms underlying the repression of systematic NLRP3 inflammasome activation remain largely unknown.

The ubiquitin-proteasome systems (UPS) play important roles in cell homeostasis, growth, and differentiation [29,30]. Protein ubiquitination is managed by three enzymes: E1-ubiquitin activating enzyme, E2-ubiquitin-conjugating enzyme, and E3-ubiquitin ligase [31]. Together with El and E2, E3 ligase catalyzes ubiquitination of a variety of proteins [32]. E3 ligases are grouped into three classes: really interesting new gene (RING), homologous to the E6-AP C terminus (HECTs), and RING between RING (RBR) [33]. Among them, Skp1-Cullin1-F-box (SCF) complex is responsible for turnover of many key proteins [34–36] and consists of three major components (CUL1, ROC1, and SKP1) along with one F-box protein [37,38]. CUL1 and ROC1 form the core of E3 ubiquitin ligase that associates with E2, whereas SKP1 links CUL1 to F-box protein [39–41]. Loss of CUL1 results in early embryonic lethal and dysregulation of Cyclin E, whereas over-expression of CUL1 is associated with cancers [43–47]. However, the role of CUL1 in regulation of NLRP3 inflammasome has not been revealed.

This study elucidates the mechanism underling repression of systematic NLRP3 inflammasome activation. Results reveal that CUL1 interacts with NLRP3 to disrupt the inflammasome assembly and also catalyzes NLRP3 ubiquitination to repress the inflammasome activation under normal physiological conditions. Moreover, CUL1 disassociates from NLRP3 to release the repression of inflammasome assembly and activation in response to inflammatory stimuli. Thus, we reveal a distinct mechanism by which CUL1 represses systematic NLRP3 inflammasome activation.

## Results

### CUL1 interacts with NLRP3 through its C-terminus

To reveal the mechanism underlying NLRP3 inflammasome repression, we screened cellular proteins interacting with NLRP3 by yeast two-hybrid analyses (Fig EV1). Several proteins, including CUL1, PGM1, and ATP1b3, were identified to interact with NLRP3 PYD (Table 1). Co-IP analyses confirmed that NLPR3 and NLRP3 PYD interacted with CUL1 but not with ATP1b3 or PGM1 (Fig 1A), and CUL11 and NLRP3 interacted with each other (Fig 1B). In macrophages, endogenous CUL1 interacted strongly with endogenous NLRP3 (Fig 1C). Immunofluorescence microscopy showed that CUL1 or NLRP3 alone diffusely distributed in the cytoplasm, whereas CUL1 and NLRP3 together were co-localized in the cells (Fig 1D).

**Figure 1.**
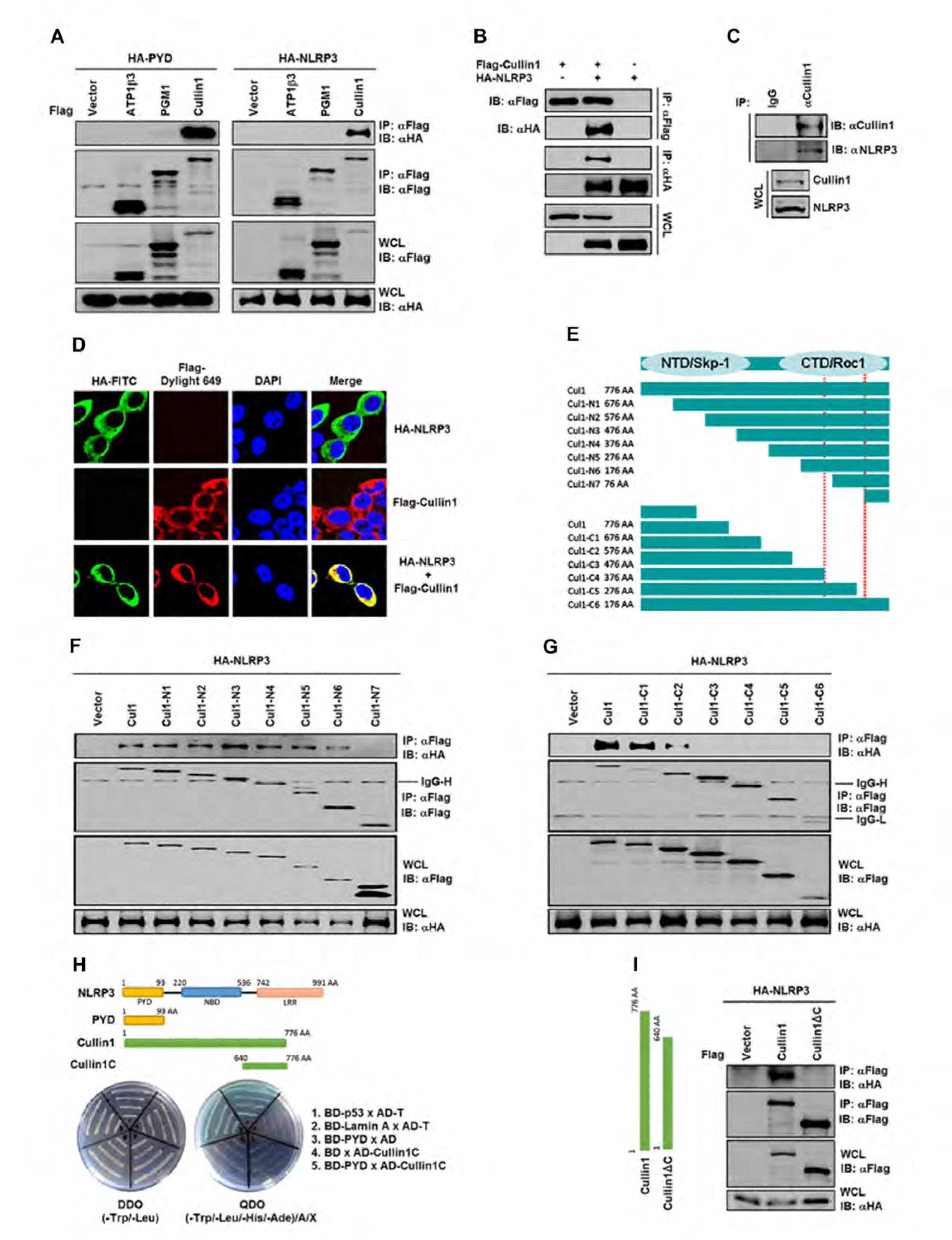
CUL1 interacts with NLRP3 through its C-terminus. **A** HEK293T cells were transfected with pCAGGS-HA-PYD or pCAGGS-HA-NLRP3 and pcDNA3.1(+)-3XFlag-ATP1β3, pcDNA3.1(+)-3XFlag-PGM1, or pcDNA3.1(+)-3XFlag-Cullin1, respectively. **B** HEK293T cells were transfected with pcDNA3.1(+)-3XFlag and pCAGGS-HA, pcDNA3.1(+)-3XFlag and pCAGGS-HA-NLRP3, or pcDNA3.1(+)-3XFlag and pCAGGS-HA-NLRP3, respectively. **C** THP-1 cells were differentiated into macrophages with PMA. **D** HEK293T cells were transfected with pcDNA3.1(+)-3XFlag-Cullin1 and pCAGGS-HA-NLRP3. The cells were immunostained with anti-Flag and anti-HA antibodies. The sub-cellular localizations of HA-NLRP3 (green), Flag-Cullin1 (Red), and nucleus marker DAPI (blue) were analyzed under confocal microscopy. **E** N terminal mutants and C terminal mutants of CUL1 were constructed and indicated. **F** HEK293T cells were transfected with pCAGGS-HA-NLRP3 and pcDNA3.1(+)-3XFlag-Cullin1 or the N terminal mutants, respectively. **G** HEK293T cells were transfected with pCAGGS-HA-NLRP3 and pcDNA3.1(+)-3XFlag-Cullin1 or the C terminal mutants, respectively. **H** Yeast strain Y2HGold was co-transformed with the combination of BD and AD plasmid, as indicated. Transfected yeast cells were grown on SD-minus Trp/Leu (DDO) plates, and colonies were replicated onto SD-minus Trp/Leu/Ade/His Plates containing Aureobasidin A and X-β-gal (QDO/A/X) to check for the expression of reporter genes. **I** HEK293T cells were transfected with pCAGGS-HA-PYD and pcDNA3.1(+)-3XFlag-Cullin1 or pcDNA3.1(+)-3XFlag-Cullin1ΔC. Data information: In (**A–C, F, G, and I**), the cell lysates were immunoprecipitated with anti-Flag, anti-HA, or anti-Cullin1 antibodies and then immunoblotted with indicated antibodies.

**Table 1.** The proteins interacted with NLRP3 as identified by yeast two-hybrid screening.

| Clone number | Comparison results after sequencing | Gene identified |
| --- | --- | --- |
| 19-1 | Correct | Cullin1 (CUL1) |
| 22-3/51-4 | Correct | Phosphoglucomutase 1 (PGM1) |
| 43-3/44-1 | Correct | ATPase Na<sup>+</sup>/K<sup>+</sup> transporting subunit beta 3 (ATP1β3) |
| 10-5 | Frameshift mutation | PNN-interacting serine/arginine-rich protein (PNISR) |
| 25-1 | Frameshift mutation | Peptidylprolyl isomerase A (PPIA) |
| 51-1 | Frameshift mutation (not in CDS) | Zinc finger protein 146 (ZNF146) |
| 55-2 | Frameshift mutation | Oxysterol binding protein-like 1A (OSBPL1A) |
| 11-6/22-5/44-6/5<br>1-3/52-3 | No significant similarity found | None |

The cellular proteins interacting with NLRP3 were identified through yeast two-hybrid screening using PYD of NLRP3 as the bait. Several cellular proteins, including Cullin1 (CUL1), phosphoglucomutase 1 (PGM1), and ATPase transporting subunit beta 3 (ATP1b3), were identified to interact with NLRP3 PYD.

To determine the region of CUL1 involved in interaction with NLRP3, a series truncated mutations of CUL1 lacking N terminus or C terminus were constructed (Fig 1E). NLRP3 interacted with CUL1, CUL1-N1, CUL1-N2, CUL1-N3, CUL1-N4, CUL1-N5, and CUL1-N6, but not with CUL1-N7 (Fig 1F), suggesting that C-terminus is involved in the interaction. NLRP3 interacted with CUL1, CUL1-C1, and CUL1-C2, but not with CUL1-C3, CUL1-C4, CUL1-C5, or CUL1-C6 (Fig 1G), demonstrating that C-terminus is required for the interaction. A CUL1 mutant (CUL1C) carrying C-terminus (aa640–aa677) and lacking N-terminus (aa1– aa639) and a NLRP3 mutant (NLRP3PYD) containing PYD (aa1–aa93) and lacking the rest part of NLRP3 (aa94–aa991) were constructed (Fig 1H, upper). Yeast two-hybrid analyses showed that CUL1C interacted with NLRP3PYD (Fig 1H, low), supporting that CUL1 C-terminus is required for interacting with NLRP3 PYD. Moreover, a CUL1 mutant (CUL1ΔC) containing only N-terminus (aa1–aa640) and lacking C-terminus (aa641–aa677) was constructed (Fig 1I, left). NLRP3 interacted with CUL1 but not with CUL1ΔC (Fig 1I, right), suggesting that C-terminus is essential for CUL1 to interact with NLRP3. Taken together, we demonstrate that CUL1 interacts with NLRP3 through its C-terminus.

### NLRP3 interacts with CUL1 by competing with ROC1

CUL1 is a scaffold protein of the SCF E3 ligase complex consisting of three components (CUL1, ROC1, and SKP-1) and one F-box protein [42], whereas NLRP3 is a sensor protein of the NLRP3 inflammasome complex consisting of three components (NLRP3, ASC, and pro-Casp-1) and the substrate (pro-IL-1β) [7]. The mutual correlations among the two complexes were evaluated. In macrophages, endogenous CUL1 interacted with NLRP3 but not Casp-1 (Fig 2A). In HEK293T cells, NLRP3 interacted with CUL1 and ROC1 but not SKP1-α or SKP1-β (Fig 2B). CUL1 interacted with ROC1, SKP1, and NLRP3 (Fig 2C, left), whereas NLRP3 interacted with CUL1 and ROC1 but not SKP-1 (Fig 2C, right). Interaction of NLRP3 with CUL1 was attenuated by ROC1, and interaction of NLRP3 with ROC1 was reduced by CUL1 (Fig 2D). Interestingly, interaction between CUL1 and NLRP3 was attenuated by ROC1 but not by SKP-1 (Fig 2E and F). However, interaction between CUL1 and ROC1 was not affected by NLRP3 (Fig 2G).

**Figure 2.**
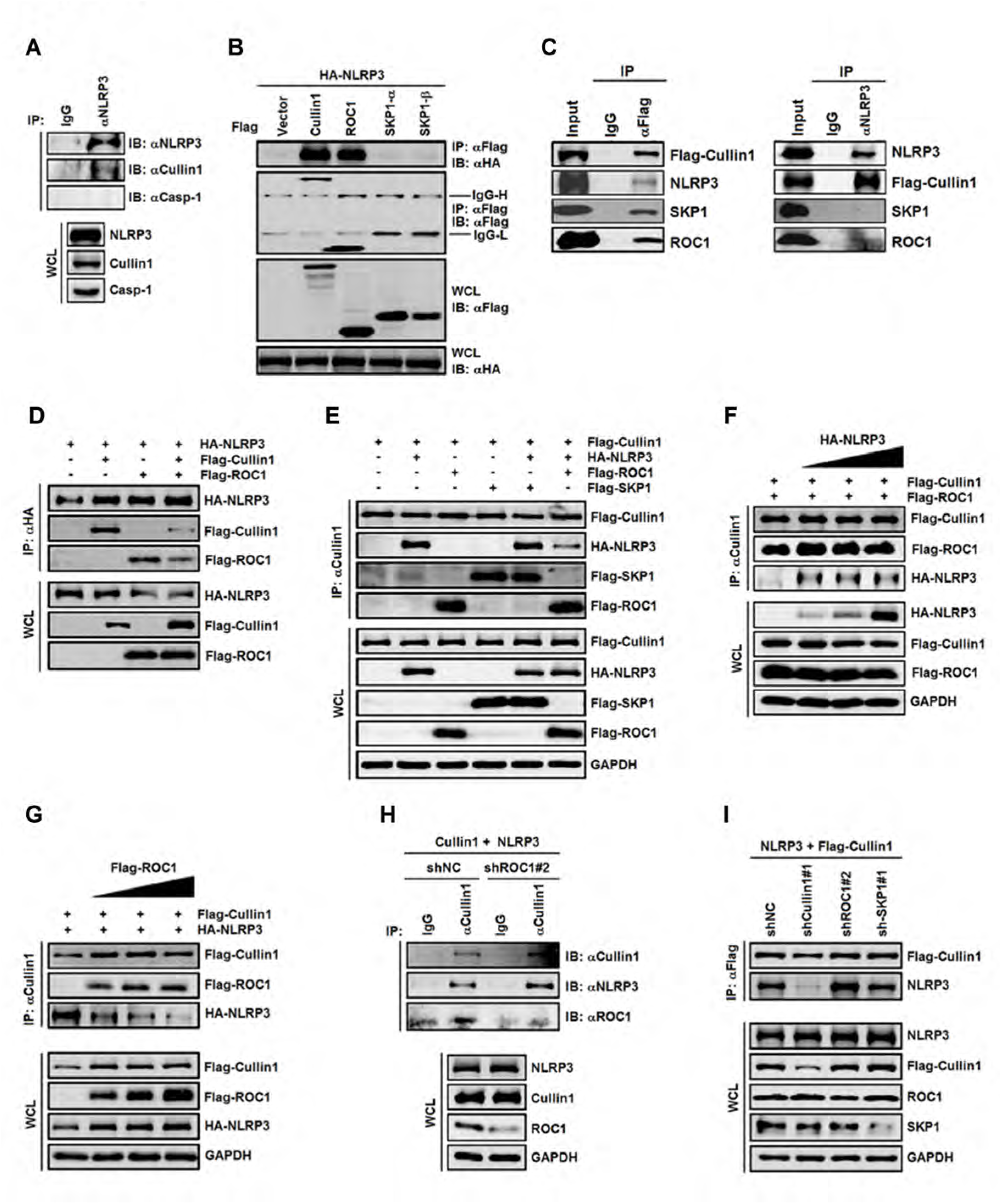
NLRP3 interacts with CUL1 by competing with ROC1. **A** The lysates of PMA-differentiated THP-1 macrophages were immunoprecipitated with anti-Cullin1 antibody and then immunoblotted with indicated antibodies. **B** HEK293T cells were transfected with pCAGGS-HA-NLRP3 and pcDNA3.1(+)-3xFlag-Cullin1, pcDNA3.1(+)-3xFlag-ROC1, pcDNA3.1(+)-3xFlag-SKP1-α and pcDNA3.1(+)-3xFlag-SKP1-β, respectively. **C** HEK293T cells were transfected with pcDNA3.1(+)-NLRP3 and pcDNA3.1(+)-3XFlag-Cullin1. **D** HEK293T cells were transfected with pCAGGS-HA-NLRP3 and pcDNA3.1(+)-3xFlag-Cullin1, pcDNA3.1(+)-3xFlag-ROC1, pcDNA3.1(+)-3xFlag-Cullin1, and pcDNA3.1(+)-3xFlag-ROC1. **E** HEK293T cells were transfected with pcDNA3.1(+)-3xFlag-Cullin1 and pCAGGS-HA-NLRP3, pcDNA3.1(+)-3xFlag-ROC1 and pcDNA3.1(+)-3xFlag-SKP1, pcDNA3.1(+)-3XFlag-SKP1 and pCAGGS-HA-NLRP3, or pcDNA3.1(+)-3xFlag-ROC1 and pCAGGS-HA-NLRP3, respectively. **F** HEK293T cells were transfected with pCAGGS-HA-NLRP3 and pcDNA3.1(+)-3xFlag-Cullin1 and pcDNA3.1(+)-3xFlag-ROC1 at different concentrations. **G** HEK293T cells were transfected with pcDNA3.1(+)-3XFlag-Cullin1 or pcDNA3.1(+)-3XFlag-ROC1 and pCAGGS-HA-NLRP3 at different concentrations. **H** HEK293T cells were transfected with pcDNA3.1(+)-3xFlag-Cullin1 and pSilencer2.1-U6-sh-Cullin1#1 or pSilencer2.1-U6-sh-Cullin1#2. **I** HEK293T cells were transfected with pcDNA3.1(+)-3xFlag-Cullin1 or pcDNA3.1(+)-3xFlag-NLRP3 and pSilencer2.1-U6-shROC1#2. Data information: In (**B–I**), the cell lysates were immunoprecipitated with anti-Flag antibody (B), anti-NLRP3 antibody (C, E), anti-HA antibody (D, F), anti-Cullin1 antibody (G–I) and then immunoblotted with indicated antibodies.

Moreover, the effects of CUL1, ROC1, and SKP1 on interaction between CUL1 and NLRP3 were evaluated using short hairpin RNAs to *Cullin-1* (sh-Cullin-1#1 and sh-Cullin-1#2), *ROC1* (sh-ROC1#1 and sh-ROC1#2), or *SKP1* (sh-SKP1#1 and sh-SKP1#2) (Fig EV2A–C). Interaction between CUL1 and NLRP3 was up-regulated by sh-ROC1#2, but interaction between CUL1 and ROC1 was down-regulated by sh-ROC1#2 (Fig 2H), suggesting that ROC1 plays an inhibitory role in interacting of CUL1 with NLRP3. Interaction between NLRP3 and CUL1 was significantly attenuated by sh-Cullin-1#1, promoted by sh-ROC1#2, but not affected by sh-SKP1#1 (Fig 2I), indicating that ROC1 represses interaction of NLRP3 with CUL1. Considering the fact that CUL1 interacts with both NLRP3 and ROC1 through its C-terminus, we speculate that NLRP3 competes with ROC1 in interacting with CUL1.

### CUL1 represses NLRP3 inflammasome in HEK293T cells

The role of CUL1 in regulation of NLRP3 inflammasome was determined in a reconstructed NLRP3 inflammasome system, in which HEK293T cells were co-transfected with plasmids encoding NLRP3 inflammasome components of (NLRP3, ASC, and pro-Casp1) and the substrate pro-IL-1β [48–51]. IL-1β secretion, IL-1β (p17) cleavage, and Casp-1 (p20) maturation were detected (Fig 3A), indicating that reconstructed NLRP3 inflammasome is active. IL-1β secretion was inhibited by CUL1 but not by ATP1b3, PGM1, or CUL1ΔC (Fig 3B and C), and IL-1β secretion, IL-1β (p17) cleavage, and Casp-1 (p20) maturation were inhibited by CUL1 but not by CUL1ΔC (Fig 3D), demonstrating that CUL1 specifically represses NLRP3 inflammasome and CUL1 C-terminus plays an essential role in the repression.

**Figure 3.**
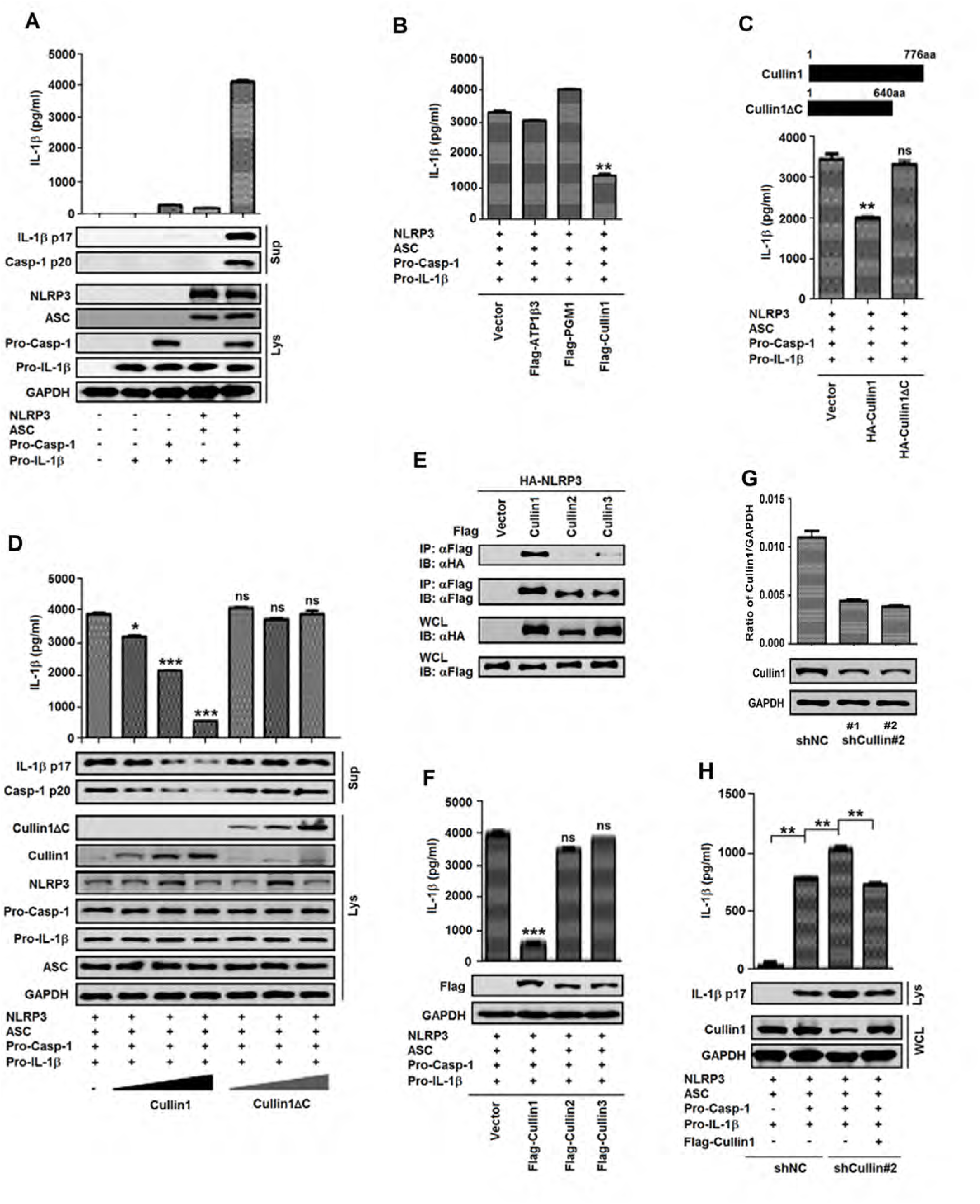
CUL1 represses NLRP3 inflammasome in HEK293T cells. **A** HEK293T cells were transfected or co-transfected with pcDNA3.1(+)-NLRP3, pcDNA3.1(+)-pro-IL-1β, pcDNA3.1(+)-pro-caspase-1, and pcDNA3.1(+)-ASC, as indicated. **B** HEK293T cells were co-transfected with pcDNA3.1(+)-NLRP3, pcDNA3.1(+)-pro-IL-1β, pcDNA3.1(+)-pro-caspase-1, and pcDNA3.1(+)-ASC, along with pcDNA3.1(+)-3Flag-ATP1β3, pcDNA3.1(+)-3Flag-PGM1, or pcDNA3.1(+)-3Flag-Cullin1, as indicated. Secreted IL-1β in the supernatants were analyzed by ELISA. **C** Daigrams of Cullin1 and Cullin1ΔC (upper). HEK293T cells were co-transfected with pcDNA3.1(+)-NLRP3, pcDNA3.1(+)-pro-IL-1β, pcDNA3.1(+)-pro-caspase-1, and pcDNA3.1(+)-ASC along with pcDNA3.1(+)-3XFlag-Cullin1 or pcDNA3.1(+)-3XFlag-Cullin1ΔC, as indicated. Secreted IL-1β in the supernatants were analyzed by ELISA. **D** HEK293T cells were co-transfected with pcDNA3.1(+)-NLRP3, pcDNA3.1(+)-pro-IL-1β, pcDNA3.1(+)-pro-caspase-1, and pcDNA3.1(+)-ASC along with pcDNA3.1(+)-3XFlag-Cullin1 or pcDNA3.1(+)-3XFlag-Cullin1ΔC. **E** HEK293T cells were co-transfected with pCAGGS-HA-NLRP3 and pcDNA3.1(+)-3XFlag-Cullin1, pcDNA3.1(+)-3XFlag-Cullin2, or pcDNA3.1(+)-3XFlag-Cullin3, respectively. The cell lysates were immunoprecipitated with anti-HA antibody and then immunoblotted with indicated antibodies. **F** HEK293T cells were co-transfected with pcDNA3.1(+)-NLRP3, pcDNA3.1(+)-ASC, pcDNA3.1(+)-pro-caspase-1, and pcDNA3.1(+)-pro-IL-1β along with pcDNA3.1(+)-3XFlag-Cullin1, pcDNA3.1(+)-3XFlag-Cullin2, or pcDNA3.1(+)-3XFlag-Cullin3, respectively. **G** HEK293T cells were transduced with lentiviruses stably expressing shRNA (sh-NC) and shRNA against Cullin1 (sh-Cullin1#1 and sh-Cullin1#2) and selected with puromycin for 2 weeks. Cullin1 and GAPDH mRNAs were determined by qRT-PCR (upper) and Cullin1 and GAPDH proteins were detected Western blot analyses (low). **H** HEK293T cells stably expressing sh-NC or sh-Cullin1#2 were co-transfected with pcDNA3.1(+)-NLRP3, pcDNA3.1(+)-pro-IL-1β, pcDNA3.1(+)-pro-caspase-1, and pcDNA3.1(+)-ASC, along with pcDNA3.1(+)-3XFlag-Cullin1. Data information: In (**A**, **D**, **F**, and **H**), secreted IL-1β in the supernatants were analyzed by ELISA (upper). Matured IL-1β (p17) and matured Casp-1 (p20) in the cell lysates were determined by immunoblot analyses with indicated antibodies (low). Data information: In (**A–D**, **G–H**), data shown are means ± SEM, *P<0.05, **P<0.01, ***P<0.0001.

CUL1 is a member in the Cullin family [41]. We determined whether other family members also regulate NLRP3 inflammasome. NLRP3 strongly interacted with CUL1 but not with CUL2 or CUL3 (Fig 3E), and IL-1β secretion was inhibited by CUL1 but not by CUL2 or CUL3 (Fig 3F), revealing that only CUL1 represses NLRP3 inflammasome. The role CUL11 in repression of NLRP3 inflammasome was further evaluated using short hairpin RNAs to *Cullin-1* (sh-Cullin-1#1 and sh-Cullin-1#2) (Fig 3G). IL-1β secretion, IL-1β (p17) cleavage, and Casp-1 (p20) maturation were up-regulated by sh-Cullin-1#2, and attenuated by CUL1 (Fig. 3H). Thus, we reveal that CUL1 represses NLRP3 inflammasome activation.

### CUL1 inhibits endogenous NLRP3 inflammasome in macrophages

The role of CUL1 in repression of endogenous NLRP3 inflammasome was determined in THP-1 differentiated macrophages. Macrophages were infected with influenza A virus (IAV) H3N2 or treated with Z-YVAD-FMK (an inhibitor of Casp-1). *IL-1β* mRNA was induced by H3N2 but not by Z-YVAD-FMK (Fig EV3A), whereas IL-1β protein was induced by H3N2 but attenuated by Z-YVAD-FMK (Fig EV3B), indicating that Casp-1 is involved in IL-1β secretion but not *IL-1β* expression. THP-1 cell lines stably expressing sh-NC, sh-NLRP3, or sh-ASC were generated (Fig EV4C and D). In these cells, IL-1β secretion was induced by H3N2 but attenuated by sh-NLRP3 or sh-ASC (Fig EV4E), suggesting NLRP3 and ASC are required for H3N2-induced secretion of IL-1β. In macrophages, IL-1β (p17) cleavage and Casp-1 (p20) maturation were induced by IAV H3N2 and lipopolysaccharides (LPS) plus adenosine triphosphate (ATP) (Fig 4A), indicating H3N2 infection induces NLRP3 inflammasome activation.

**Figure 4.**
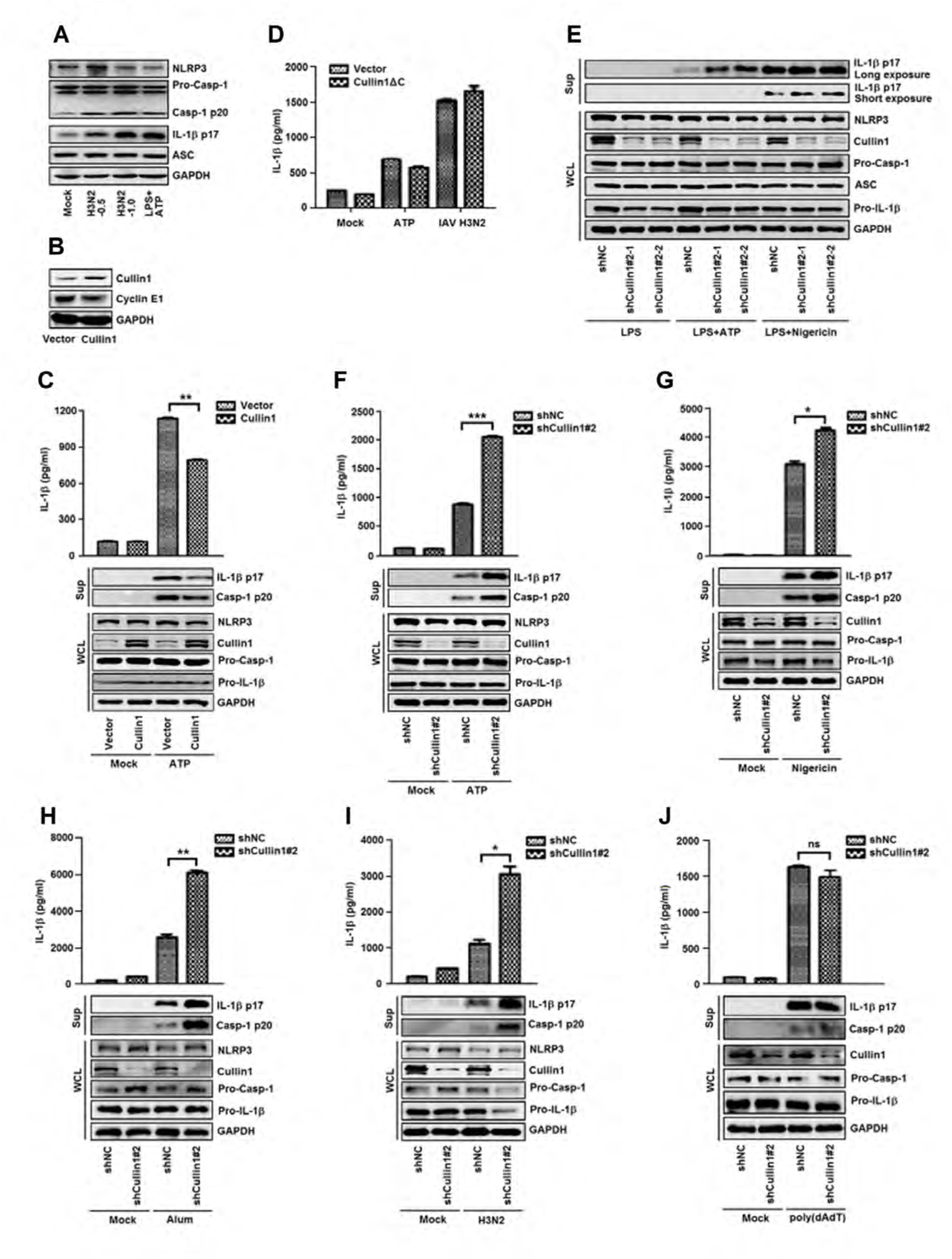
CUL1 inhibits endogenous NLRP3 inflammasome in macrophages. **A** PMA-differentiated THP-1 macrophages were infected with H3N2 for 24 h or treated with LPS and ATP for 30 min. The cell lysates were immunoblotted with indicated antibodies. **B** CUL1, Cyclin E1, and GAPDH proteins were examined by Western blot analysis in THP-1 cells stably expressing CUL1. **C** Macrophages stably expressing CUL1 were treated with ATP for 30 min. **D** Macrophages stably expressing CUL1ΔC were treated with ATP or infected with H3N2. Secreted IL-1β in the supernatants were analyzed by ELISA. **E** Macrophages stably expressing sh-Cullin1#2 were treated with LPS for 6 h, LPS for 6 h and ATP for 30 min, or LPS for 6 h and Nigericin for 1 h, respectively. Matured IL-1β (p17) in the cell lysates was determined by immunoblot analysis with indicated antibodies. **F** Macrophages stably expressing sh-Cullin1#2 were treated with ATP for 30 min. **G** Macrophages stably expressing sh-Cullin1#2 were treated with Nigericin for 1 h. **H** Macrophages stably expressing sh-Cullin1#2 were treated with Alum for 6 h **I** Macrophages stably expressing sh-Cullin1#2 were infected with IAV H3N2 for 24 h **J** Macrophages stably expressing sh-Cullin1#2 were treated with Poly(dA:dT)/LyoVec^TM^ for 12 h. Data information: In (**C**, **F–J**), secreted IL-1β in the supernatants were analyzed by ELISA (upper). Matured IL-1β (p17) and matured Casp-1 (p20) in the cell lysates were determined by immunoblot analysis with indicated antibodies (low). Data information: In (**C**, **D**, **F–J**), data shown are means ± SEM, *P<0.05, **P<0.01, ***P<0.0001.

To evaluate the role of CUL1 in regulation of NLRP3 inflammasome, we generated a THP-1 cell line stably expressing *Cullin1* mRNA and CUL1 protein (Fig EV3F). Cyclin E1 was down-regulated by CUL1 (Fig 4B), which is consistent with previous reports [52,53]. The stable cells were differentiated into macrophages and then stimulated with ATP. ATP-induced IL-1β secretion, IL-1β (p17) cleavage, and Casp-1 (p20) maturation were attenuated by CUL1 (Fig 4C). A THP-1 cell line stably expressing CUL1ΔC was also established (Fig EV3G). The cells were differentiated into macrophages and then treated with ATP or infected with H3N2. IL-1β secretion was induced by ATP or H3N2 but not by CUL1ΔC (Fig 4D). These results demonstrate that CUL1 specifically represses NLRP3 inflammasome activation. The role of CUL1 in regulation of NLRP3 inflammasome was further evaluated in THP-1 cells stably expressing sh-Cullin-1#1 or sh-Cullin-1#2 (Fig EV3H). Cyclin E1 was up-regulated by sh-Cullin-1#2 (Fig EV3I), indicating that they are effective. The cells were differentiated into macrophages and then stimulated with LPS, LPS+ATP, or LPS+Nigericin. IL-1β (p17) cleavage and Casp-1 (p20) maturation were induced by LPS, LPS+ATP, or LPS+Nigericin, and further enhanced by sh-Cullin-1#2 (Fig 4E), indicating that knock-down of *Cullin-1* facilitates NLRP3 inflammasome activation.

NLRP3 inflammasome activation is triggered by diverse stimuli or agonists [54,55]. IL-1β secretion, IL-1β (p17) cleavage, and Casp-1 (p20) maturation were induced by ATP, Nigericin, Alum, and H3N2, and such inductions were further up-regulated by sh-Cullin-1#2 in macrophages (Fig 4F–I), suggesting that knock-down of Cullin-1 up-regulates NLRP3 inflammasome. However, IL-1β secretion, IL-1β (p17) cleavage, and Casp-1 (p20) maturation were induced by poly(dAdT) but such induction was not affected by sh-Cullin-1#2 (Fig 4J), indicating that CUL1 has no effect on AIM2 inflammasome. Taken together, we demonstrate that CUL1 specifically represses NLRP3 inflammasome activation in macrophages.

### CUL1 interacts with NLRP3 to repress inflammasome assembly

The mechanism by which CUL1 represses NLRP3 inflammasome was elucidated. NLRP3 inflammasome is a multi-protein platform that comprises NLRP3, ASC, and Casp-1 [55]. We revealed that ASC strongly interacted with NLRP3 PYD in 293T cells (Fig EV4). Interestingly, CUL1 interacted with NLRP3 and its three domains (PYD, NBD, and LRR) (Fig 5A) but not with ASC or Casp-1 (Fig 5B), suggesting that CUL1 is specifically and tightly associated with NLRP3.

**Figure 5.**
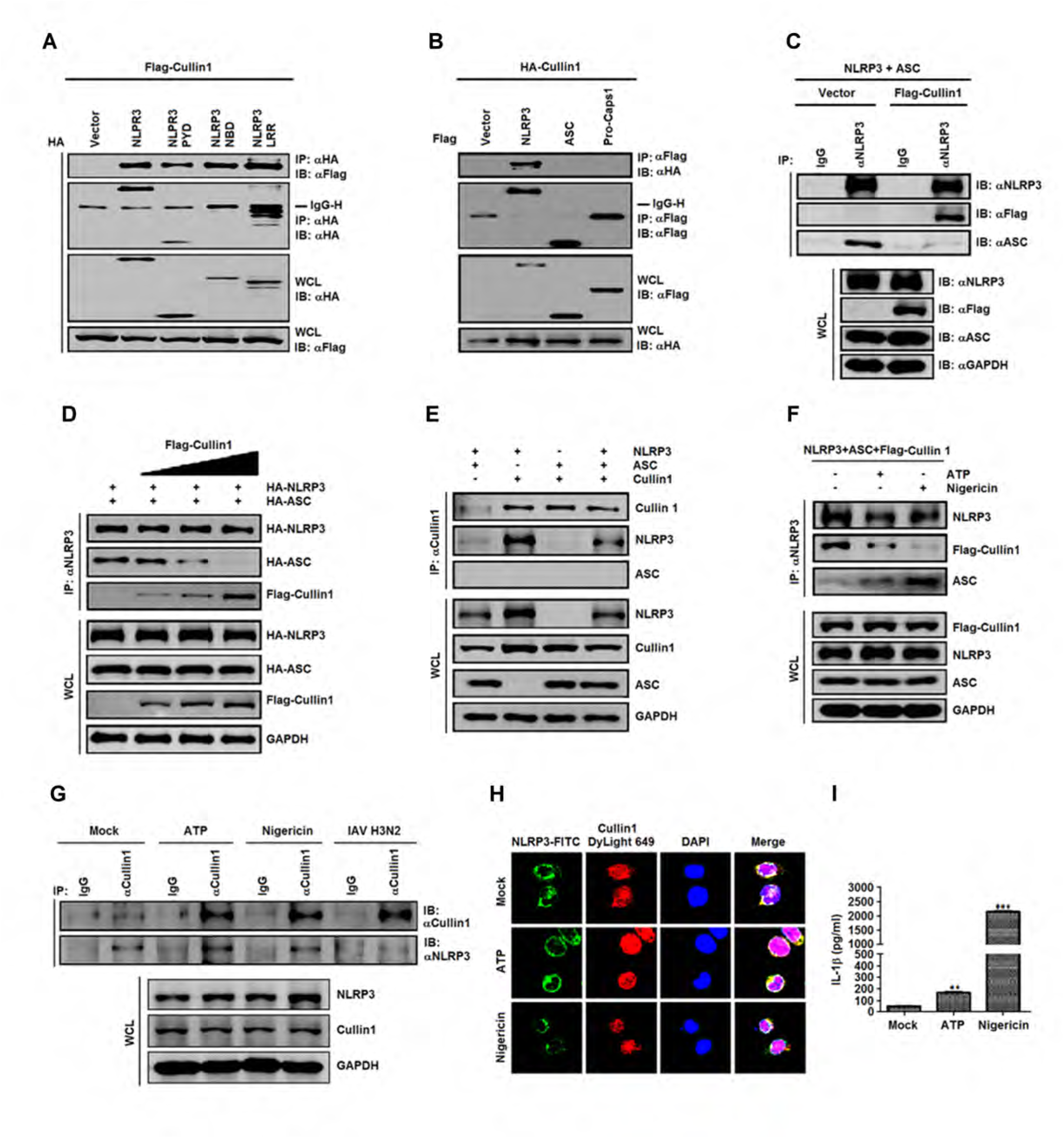
CUL1 interacts with NLRP3 to repress inflammasome assembly. **A** HEK293T cells were transfected with pcDNA3.1(+)-3XFlag-Cullin1 and pCAGGS-HA-NLRP3, pCAGGS-HA-PYD, pCAGGS-HA-NBD, pCAGGS-HA-LLR, respectively. **B** HEK293T cells were transfected with pCAGGS-HA-Cullin1 and pcDNA3.1(+)-3XFlag-NLRP3, pcDNA3.1(+)-3XFlag-ASC, or pcDNA3.1(+)-3XFlag-pro-caspase-1, respectively. **C** HEK293T cells were transfected with pcDNA3.1(+)-NLRP3 and pcDNA3.1(+)-ASC along with pcDNA3.1(+)-3XFlag-Cullin1. **D** HEK293T cells were transfected with pCAGGS-HA-NLRP3 and pCAGGS-HA-ASC along with pcDNA3.1(+)-3XFlag-Cullin1 at different concentrations. **E** HEK293T cells were co-transfected with pcDNA3.1(+)-NLRP3 and/or pcDNA3.1(+)-ASC along with pcDNA3.1(+)-3XFlag-Cullin1, as indicated. **F** HEK293T cells were co-transfected with pcDNA3.1(+)-NLRP3, pcDNA3.1(+)-ASC, pcDNA3.1(+)-3XFlag-Cullin1 and then treated with 5 mM ATP for 30 min and 10 μM Nigericin for 1 h. **G** PMA-differentiated THP-1 macrophages were stimulated with ATP or Nigericin, and infected with H3N2. **H** PMA-differentiated THP-1 macrophages were stimulated with ATP or Nigericin. The cells were immunostained with anti-NLRP3 and anti-Cullin1 antibodies. The sub-cellular localizations of endogenous NLRP3 (green), endogenous Cullin1 (Red), and nucleus marker DAPI (blue) were analyzed under confocal microscopy. **I** PMA-differentiated THP-1 macrophages were stimulated with ATP or Nigericin. The cells were immunostained with anti-NLRP3 and anti-Cullin1 antibodies. The supernatants were analyzed by ELISA for IL-1β secretion. Data shown are means ± SEM, *p<0.05, **p<0.01, ***p<0.0001. Data information: In (**A–G**), the cell lysates were immunoprecipitated with anti-HA antiboody (A), anti-Flag antibody (B, C), anti-NLRP3 antibody (D, F), or anti-Cullin1 antibody (E, G) and then immunoblotted with indicated antibodies.

The effect of CUL1 on interaction between NLRP3 and ASC was evaluated. ASC interacted with NLRP3 in the absence of CUL1 but failed to interact with NLRP3 in the presence of CUL1 (Fig 5C), and interaction of ASC with NLRP3 was attenuated by CUL1 (Fig 5D). Because CUL1 interacts with NLRP3 PYD [16–19], it is possible that CUL1 may compete with ASC PYD in interacting with NLRP3 PYD, and thus disrupt NLRP3-ASC complex assembly. Interestingly, interaction between NLRP3 and CUL1 was attenuated by ASC (Fig 5E), interaction between NLRP3 and ASC was stimulated by ATP or Nigericin, and interaction between NLRP3 and CUL1 was inhibited by ATP or Nigericin (Fig 5F). These results reveal that CUL1 inhibits NLRP3 inflammasome through competing with ASC, leading to disruption of inflammasome assembly.

Moreover, in THP-1 differentiated macrophages, interaction between endogenous CUL1 and NLRP3 was significantly attenuated by ATP, Nigericin, or H3N2 (Fig 5G), indicating that CUL1 disassociates from NLRP3 in response to stimuli. Confocal microscopy analyses showed that in the absence of stimuli, endogenous NLRP3 and CUL1 were co-localized and mainly distributed in the cytoplasm of macrophages, however, in the presence of ATP and Nigericin, endogenous NLRP3 formed spots in the cytoplasm and CUL1 was mainly located in the nucleus (Fig 5H), confirming that CUL1 is disassociated for NLRP3 during induction of inflammasome activation. IL-1β secretion was induced by ATP and significantly stimulated by Nigericin (Fig 5I), suggesting that NLRP3 inflammasome is activated. Taken together, we reveal that CUL1 interacts with NLRP3 to block the binding of ASC to NLRP3 and thereby repress NLRP3 inflammasome assembly in resting macrophages, whereas CUL1 disassociates from NLRP3 to allow ASC to bind NLRP3 and thereby release the repression of NLRP3 inflammasome assembly in response to stimuli.

### CUL1 promotes NLRP3 ubiquitination but not protein degradation

Ubiquitination of NLRP3 is essential for NLRP3 inflammasome repression [57], whereas deubiquitination of NLRP3 is critical for inflammasome activation [58,59]. CUL1 is a component of the SCF E3 ubiquitin ligase that catalyzes ubiquitination of a variety of proteins [32]. Thus, we investigated that the role of CUL1 in regulation of NLRP3 ubiquitination. NLRP3 ubiquitination increased as the concentration of ubiquitin increased (Fig 6A), indicating that NLRP3 is ubiquitinated under these conditions. NLRP3 ubiquitination was significantly promoted by CUL1 but not by ROC1 (Fig 6B and C), demonstrating that CUL1 catalyzes NLRP3 ubiquitination. By using UbPred software [60] to predicate potential ubiquitination sites in NLRP3, we revealed that one Lys (Lys689) is high confidence and six Lys (Lys93, Lys192, Lys194, Lys324, Lys430, and Lys696) are medium confidence. Thus, we constructed seven mutations of CUL1, in which the K residues were replaced by R residues (Fig 6D). Like CUL1, the mutants K93R, K192R, K194R, K324R, K430R, and K696R facilitated NLRP3 ubiquitination, but K689R failed to act (Fig 6E), confirming that Lys689 is a significant NLRP3 ubiquitination acceptor site.

**Figure 6.**
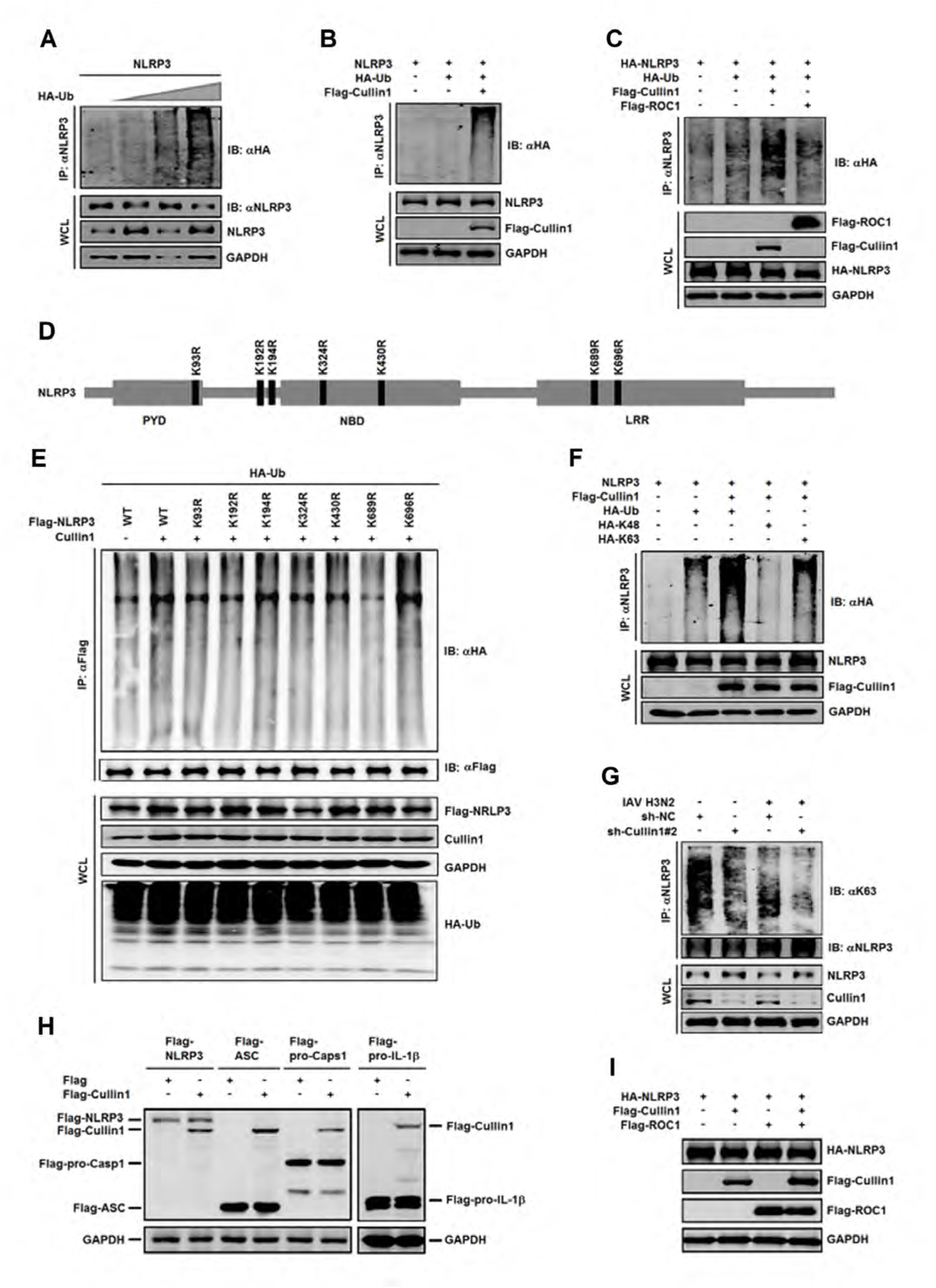
CUL1 promotes NLRP3 ubiquitination but not protein degradation. **A** HEK293T cells were transfected with pcDNA3.1(+)-NLRP3 and pCAGGS-HA-Ub at different concentrations. **B** HEK293T cells were transfected with pcDNA3.1(+)-NLRP3, pcDNA3.1(+)-HA-Ub, and/or pcDNA3.1(+)-3Flag-Cullin1, respectively. **C** HEK293T cells were transfected with pcDNA3.1(+)-NLRP3, pcDNA3.1(+)-HA-Ub, pcDNA3.1(+)-3Flag-Cullin1 or pcDNA3.1(+)-3Flag-ROC1, as indicated. **D** Daigrams of full-length NLRP3 and its point mutants. The potential ubiquitination sites in NLRP3 among the Lys residues were predicated by using UbPred software. One Lys (Lys689) is high confidence and six Lys (Lys93, Lys192, Lys194, Lys324, Lys430, and Lys696) are medium confidence. Seven mutations of CUL1 (K93R, K192R, K194R, K324R, K430R, K689R, and K696R), in which the K residues were replaced by R residues. **E** HEK293T cells were co-transfected with pcDNA3.1(+)-HA-Ub and pcDNA3.1(+)-Cullin1, along with pcDNA3.1(+)-3xFlag-NLRP3 or its mutants, respectively. **F** HEK293T cells were co-transfected with pcDNA3.1(+)-NLRP3 and/or pcDNA3.1(+)-HA-Ub, pcDNA3.1(+)-3Flag-Cullin1, pcDNA3.1(+)-HA-K48 or pcDNA3.1(+)-HA-K63, as indicated. **G** PMA-differentiated THP-1 macrophages stably expressing sh-NC and sh-Cullin1#2 were infected with IAV H3N2. **H** HEK293T cells were transfected with pcDNA3.1(+)-3XFlag-NLRP3, pcDNA3.1(+)-3XFlag-ASC, pcDNA3.1(+)-3XFlag-pro-caspase-1, pcDNA3.1(+)-3XFlag-pro-IL-1β, or pcDNA3.1(+)-3xFlag-Cullin1, as indicated. **I** HEK293T cells were transfected with pCAGGS-HA-NLRP3 and/or pcDNA3.1(+)-3XFlag-Cullin1, or pcDNA3.1(+)-3XFlag-ROC1. Data information: In (**A–C**), the lysates were immunoprecipitated with anti-NLRP3 antibody and immunoblotted with indicated antibodies. In (**E–I**), the lysates were immunoprecipitated with anti-Flag antibody (E) or anti-NLRP3 antibody (F, G) and then immunoblotted with indicated antibodies. The lysates were immunoblotted with the indicated antibodies (H, I).

NLRP3 is ubiquitinated with both K48 and K63 linkages in macrophages [57]. NLRP3 ubiquitination was detected in the presence of Ub and significantly facilitated by CUL1, and CUL1 promoted NLRP3 ubiquitination in the presence of K63 but not in the presence of K48 (Fig 6F), indicating that CUL1 catalyzes K68-linked ubiquitination of NLRP3. THP-1 differentiated macrophages stably expressing sh-NC or sh-Cullin-1#2 were infected with H3N2. K63-linked ubiquitination of endogenous NLRP3 was down-regulated by sh-Cullin-1#2 and H3N2, and H3N2-mediated reduction of NLRP3 ubiquitination was further attenuated by sh-Cullin-1#2 (Fig 6G), confirming that CUL1 promotes K68-linked ubiquitination of endogenous NLRP3.

For our surprise, the levels of NLRP3, ASC, pro-Casp-1, and pro-IL-1β proteins were not affected by CUL1 (Fig 6H and Fig EV5), suggesting that CUL1 regulates NLRP3 inflammasome independent of protein degradation. NLRP3 level was not affected by CUL1 or ROC1 or both proteins (Fig 6I), demonstrating that CUL1 and ROC1 have no effect on NLRP3 stability. We noticed that this result is different from previous reports showing that dopamine and TRIM31 inhibit NLRP3 inflammasome by promoting proteasomal degradation of NLRP3 [27].

### CUL1 suppresses IL-1β activation and inflammatory response in mice

The effect of CUL1 on peritoneal inflammation was determined in a mouse peritonitis model described previously [61]. Three siRNAs targeting to mouse *Cullin-1* (siR-Cullin1#1, siR-Cullin1#2, and siR-Cullin1#3) were generated. In mice L929 cells, *Cullin-1* mRNA and CUL1 protein were significantly down-regulated by siR-Cullin1#1, reduced by siR-Cullin1#2, and relatively unaffected by siR-Cullin1#3 (Fig 7A). C57BL/6 mice were injected with siR-Cullin1#1 and then stimulated with Alum to trigger peritonitis. Endogenous CUL1 in mice peritoneal exudates cells (PECs) was down-regulated by siR-Cullin1#1, and secreted IL-1β protein in mice sera was significantly up-regulated by siR-Cullin1#1 (Fig 7B). Secreted IL-1β in mice peritoneal lavage fluid was induced by Alum and further enhanced by siR-Cullin1#1 (Fig 7C), however, secreted IL-6 was induced by Alum but relatively unaffected by siR-Cullin1#1 (Fig 7D). The total numbers of PECs (Fig 7E) and Neutrophils (Fig 7F) in peritoneal cavity were stimulated by Alum and further enhanced by siR-Cullin1#1. Moreover, inflammatory response in mice spleen was slightly induced by Alum, but significantly facilitated by siR-Cullin1#1 (Fig 7G). The findings demonstrate that CUL1 suppress NLRP3 inflammasome activation and represses inflammatory response in mice. Taken together, we reveal a distinct mechanism by which CUL1 represses systemic NLRP3 inflammasome activation (Fig 8).

**Figure 7.**
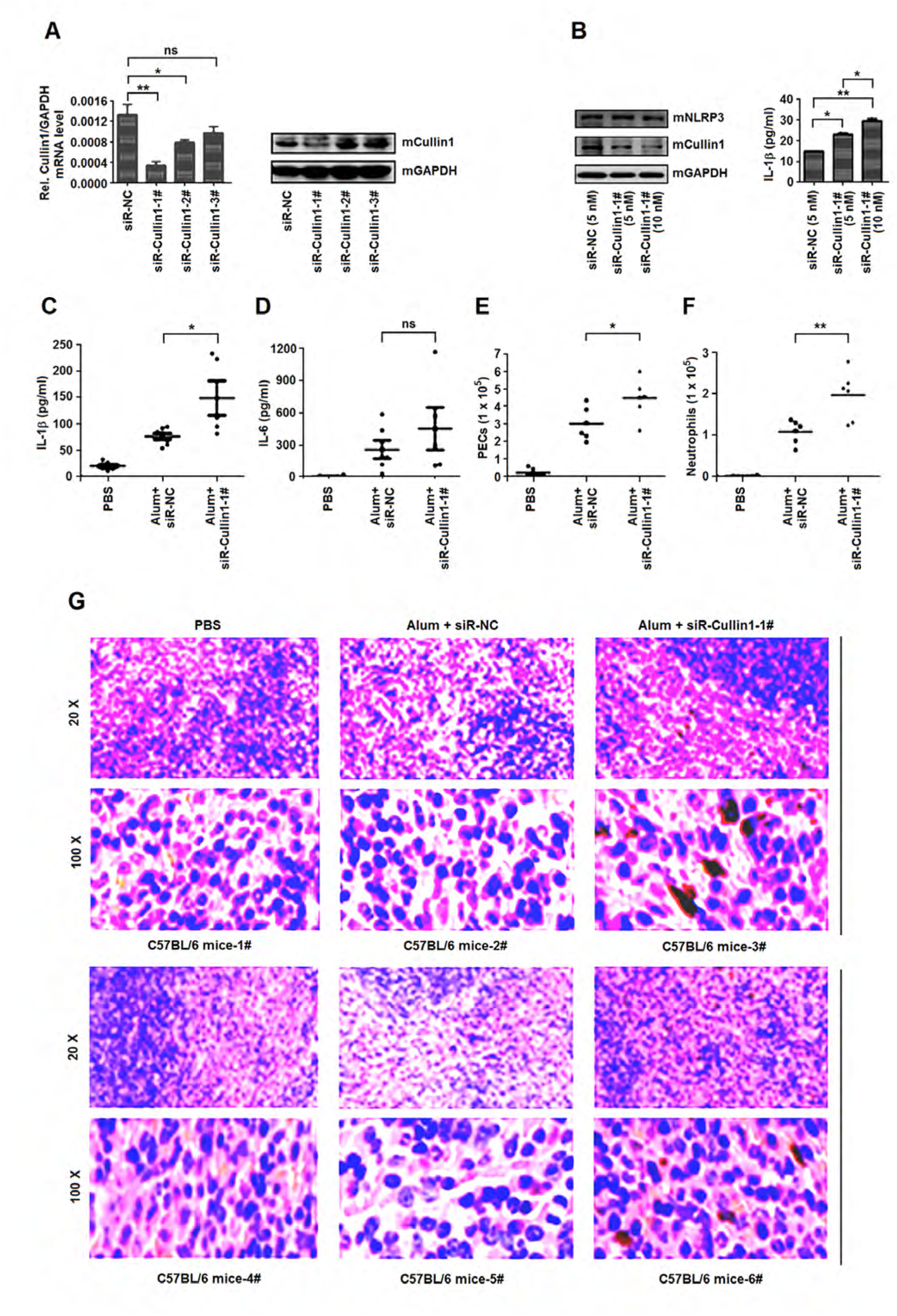
CUL1 suppresses IL-1β secretion and inflammatory response in mice. **A** L929 cells were transfected with siR-NC (25 nM), siR-Cullin1#1 (25 nM), siR-Cullin1#2 (25 nM), or siR-Cullin1#3 (25 nM). *Cullin1* and *GAPDH* mRNAs were determined by qRT-PCR (left) and CUL1 and GAPDH proteins were detected Western blot analyses (right). **B** C57BL/6 mice were injected with siR-NC (5 nM), siR-Cullin1#1 (5 nM), or siR-Cullin1#1 (10 nM) for 60 h, and the peritonitis of treated mice were then injected with Alum for 12 h. The endogenous mCUL1, mNLRP3, and mGAPDH proteins in the mice peritoneal exudates cells (PECs) were detected by Western blot analyses (left). Secreted IL-1β in the mice sera were analyzed by ELISA (right). **C** C57BL/6 mice (six mice per group) were injected with siR-NC or siR-Cullin1#1 for 60 h, and the peritonitis of treated mice were then injected with Alum. Secreted IL-1β in mice peritoneal lavage fluid were analyzed by ELISA. **D** C57BL/6 mice (six mice per group) were injected with siR-NC or siR-Cullin1#1 for 60 h, and the peritonitis of treated mice were then injected with Alum. Secreted IL-6 in mice peritoneal lavage fluid was analyzed by ELISA. **E** C57BL/6 mice (six mice per group) were injected with siR-NC or siR-Cullin1#1 for 60 h, and the peritonitis of treated mice were then injected with Alum. The total numbers of PECs in peritoneal cavity were stimulated. **F** C57BL/6 mice (six mice per group) were injected with siR-NC or siR-Cullin1#1 for 60 h, and the peritonitis of treated mice were then injected with Alum. The total numbers of Neutrophils (F) in peritoneal cavity were stimulated. **G** C57BL/6 mice (six mice per group) were injected with siR-NC or siR-Cullin1#1 for 60 h, and the peritonitis of treated mice were then injected with Alum. Histopathological changes in the mice spleen tissues were examined by H&E staining. Data information: In (**A–F**), data shown are means ± SEM, *P<0.05, **P<0.01, ***P<0.0001.

**Figure 8.**
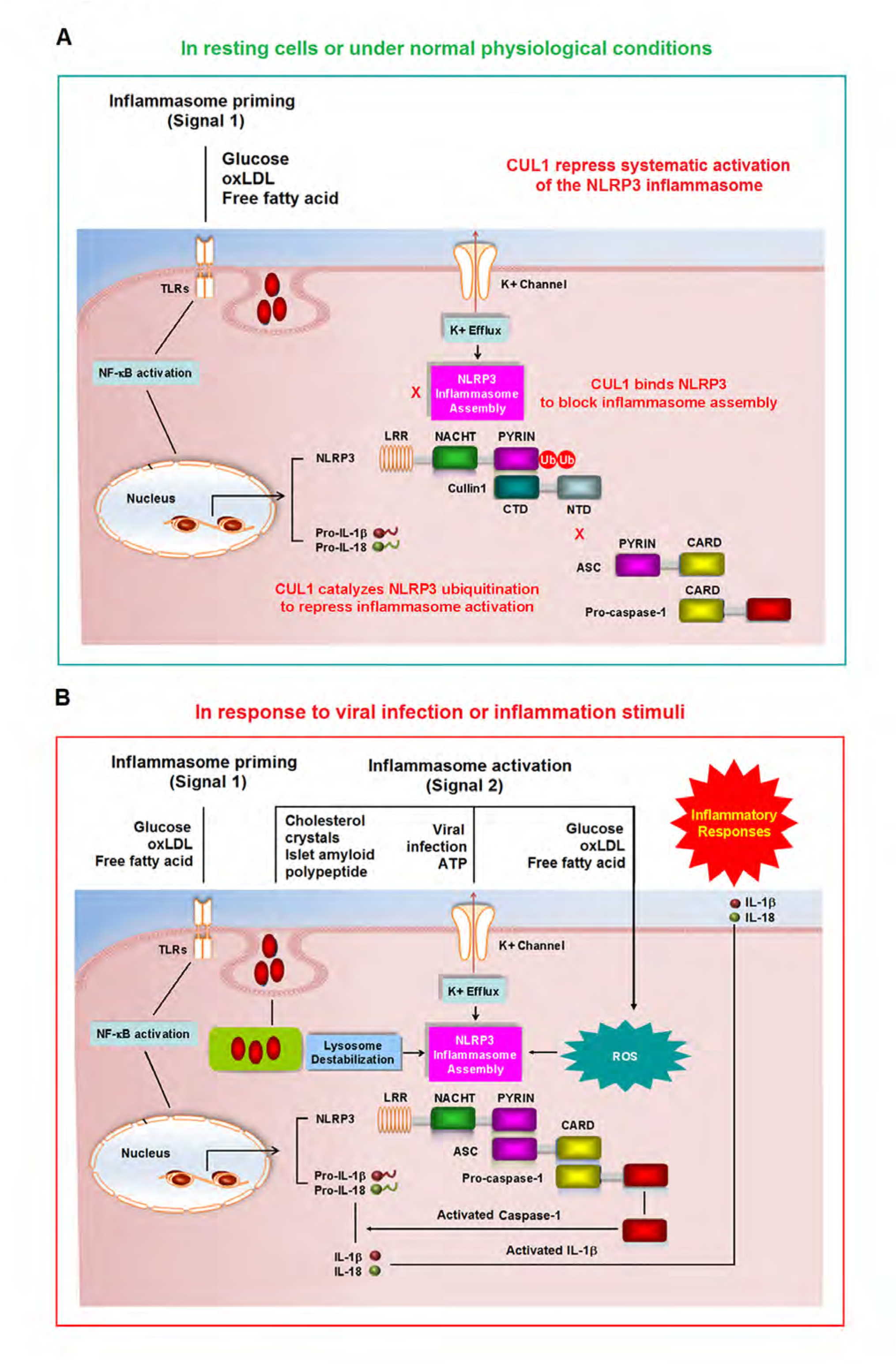
A proposed mechanism underlying CUL1-mediated repression of systematic NLRP3 inflammasome activation. **A** In resting cells and under normal physiological conditions, CUL1 interacts with NLRP3 to disrupt the NLRP3 inflammasome assembly, and catalyzes NLRP3 ubiquitination to repress the NLRP3 inflammasome activation. **B** In response to and inflammasome stimuli and pathogen infection, CUL1 disassociates from NLRP3 to release the repression of NLRP3 inflammasome assembly and activation. ASC binds NLRP3 to form ASC oligomers, which provides a platform for activation of pro-Casp-1 that regulates the maturation and secretion of IL-1β and IL-18, which in turn initiate multiple signaling pathways and drive inflammatory responses.

## Discussion

The NLRP3 inflammasome is activated in response to pathogen-associated molecular patterns (PAMP) and danger-associated molecular patterns (DAMP) to induce inflammatory responses [62,63]. Precise and tight control of NLRP3 inflammasome is critical to adequate immune protection and limit detrimental responses. Although significant attempts have been made and negative regulations are employed by hosts and pathogens to inhibit NLRP3 inflammasome activation [64], the mechanisms underlying repression of systematic NLRP3 inflammasome activation remain largely unknown.

This study initially reveals that CUL1 interacts with NLRP3 PYD through its C-terminus, which is required for nuclear localization and SCF ubiquitin ligase assembly [65]. CUL1 interacts with all three domains (PYD, NACHT, and LRR) of NLRP3, and endogenous NLRP3 and CUL1 co-localized and distributed in the cytoplasm of macrophages. Thus, we demonstrate that CUL1 strongly associates with NLRP3. CUL1, a scaffold protein of SCF E3 ligase complex that consists three major components (CUL1, ROC1, and SKP-1) [42], and NLRP3, is a sensor protein of the NLRP3 inflammasome complex that also contains three major components (NLRP3, ASC, and pro-Casp-1) [7,56]. The mutual correlations among the two complexes are revealed: CUL1 interacts with NLRP3 but not pro-Casp-1 or ASC, whereas NLRP3 interacts with CUL1 and ROC1 but not SKP-1. Interaction of NLRP3 with CUL1 is attenuated by ROC1, interaction of NLRP3 with ROC1 is reduced by CUL1, and interaction between CUL1 and ROC1 is not affected by NLRP3. Considering the fact that CUL1 interacts with both NLRP3 and ROC1, we propose that NLRP3 competes with ROC1 in interacting with CUL1. NLRP3 binds MAVS, which is critical for activating NLRP3 inflammasome [66,67]. We speculate that binding between CUL1 and NLRP3 may play a role in regulating MAVS, although it needs to be further investigated. Nevertheless, the results suggest that SCF E3 ligase and NLRP3 inflammasome are highly associated, which may play roles in controlling important cellular functions.

Interaction between NLRP3 and ASC is necessary for inflammasome assembly, which provides a platform for pro-Casp-1 activation [16,18,19]. Interaction between NLRP3 and ASC is repressed by CUL1, indicating CUL1 disrupts inflammasome assembly. NLRP3 inflammasome is activated by LPS, ATP, Nigericin, Alum, and H3N2, but such induction is attenuated by CUL1 and enhanced by sh-Cullin-1, revealing that CUL1 represses NLRP3 inflammasome activation. Moreover, by using a mouse peritonitis mode, we further reveal that knock-down of Cullin-1 up-regulates IL-1β secretion in mice sera and peritoneal lavage fluid, increases PECs and Neutrophils in peritoneal cavity, and induces inflammatory responses in mice spleen, demonstrating that CUL1 suppress NLRP3 inflammasome and inflammatory response. Taken together, we demonstrate that CUL1 represses NLRP3 inflammasome assembly and activation.

Ubiquitination of NLRP3 is essential for NLRP3 inflammasome repression [57,68], whereas deubiquitination of NLRP3 is critical for the inflammasome activation [23,58]. This study demonstrates that CUL1 catalyzes K68-linked ubiquitination of NLRP3, and CUL1 represses NLRP3 inflammasome activation through catalyzing ubiquitination. However, the levels of NLRP3, ASC, and pro-Casp-1 are not affected by CUL1, indicating that CUL1 represses NLRP3 inflammasome independent of protein degradation. This result is different from previous reports showing that TRIM31 inhibits NLRP3 inflammasome by promoting proteasomal degradation of NLRP3 [27,57], and FBXL3 interacts with NLRP3 to facilitate SCFFBXL2-mediated degradation of NLRP3 [68]. Thus, CUL1, as an important component of the SCF E3 ubiquitin ligase, represses NLRP3 inflammasome through a distinct mechanism: catalyzes NLRP3 ubiquitination and has no effect on protein stability. This unique mechanism ensures to keep NLRP3 in check by restricting its function without destroying the protein.

Interestingly, interaction between NLRP3 and CUL1 is inhibited by ATP or Nigericin and attenuated by ASC. Because CUL1 interacts with NLRP3 PYD, and thus we propose that CUL1 inhibits NLRP3 inflammasome through competing with ASC in interacting with NLRP3 to disrupt inflammasome assembly. In addition, interaction between endogenous CUL1 and NLRP3 in macrophages is attenuated by ATP, Nigericin, and H3N2. Moreover, in the presence of ATP or Nigericin, endogenous NLRP3 forms small spot in cytoplasm and CUL1 mainly distributes in the nucleus. Taken together, these results demonstrate that CUL1 disassociates for NLRP3 to release NLRP3 inflammasome repression in response to inflammatory stimuli and pathogen infections.

In conclusion, we reveal a distinct mechanism by which CUL1 represses systemic NLRP3 inflammasome activation. In resting cells and under normal physiological conditions, CUL1 interacts with NLRP3 to disrupt the inflammasome assembly, and catalyzes NLRP3 ubiquitination to repress the inflammasome activation (Fig. 8A). However, in response to inflammatory stimuli, CUL1 disassociates from NLRP3 to release the repression of inflammasome assembly and activation (Fig. 8B).

## Materials and Methods

### Animal study

Female C57BL/6 mice (6 weeks) were obtained from Hubei Provincial Center for Disease Control and Prevention (Wuhan, China). All animal experiments were undertaken in accordance with the National Institutes of Health Guide for the Care and Use of Laboratory Animals, with the approval of the Wuhan University animal care and use committee guidelines.

Mice peritoneal exudates cells (PECs) were harvested and Red Blood Cell (RBC) Lyses Buffer was added to PECs. The cells were washed twice with PBS buffer supplemented with 2% BSA. For PECs, the cells were directly analyzed by flow cytometry (Beckman Coulter).

For neutrophils, PECs were Fc-blocked by Trustain fcX^TM^ anti-mouse CD16/32 Antibody for 10 minutes prior to staining, then PECs in PBS buffer supplemented with 2% BSA are incubated with Anti-FITC-CD11b and Anti-APC-Ly-6G on ice for 1h in the dark. Cells were washed twice with PBS buffer supplemented with 2% BSA, and analyzed by flow cytometry (Beckman Coulter).

All spleen tissues were fixed in 4% paraformaldehyde and Histopathological changes in the spleen tissue were examined by H&E staining.

For induction of peritonitis, mice were injected with 700 μg Alum (Thermo scientific). For analysis of inflammatory cell subsets and cytokines in the peritoneal cavity, mice were killed 8h after alum injection and peritoneal cavities were washed with 5 ml of PBS. PECs and neutrophils were collected and analyzed by flow cytometry and the peritoneal fluids were concentrated for ELISA analysis.

All mouse siRNAs were synthesized as following: siR-Cullin1#1: 5’-GCTTGTGGTCGCTTCATAA-3’; siR-Cullin1#2: 5’-GGTTGTATCAACTGTCCAA-3’; siR-Cullin1#3: 5’-GGTCGCTTCATAAACAACA-3’. Mice were intraperitoneally injected with siCullin1-1 and negative control for 60–72 h and all these siRNAs were specially modified by 5’ Cho1 and 2’ OMe. All siRNA were synthesized by RiboBio (Guangzhou, China).

### Cell lines and cultures

Human embryonic kidney cell line (HEK293T) was purchased from American Type Culture Collection (ATCC) (Manassas, VA, USA). Human monocytic cell line (THP-1) was a gift from Dr. Bing Sun of Institute of Biochemistry and Cell Biology, Shanghai Institute for Biological Sciences. HEK293T cells were maintained in DMEM purchased from Gibco (Grand Island, NY, USA) supplemented with 10% fetal bovine serum (FBS), 100 U/ml penicillin, and 100 μg/ml streptomycin sulfate. THP-1 cells were maintained in RPMI 1640 medium supplemented with 10% FBS, 100 U/ml penicillin, and 100 μg/ml streptomycin sulfate. THP-1 cells were differentiated for 12 h with 100 ng phorbol-12-myristate-13-acetate (PMA).

### Reagents

Lipopolysaccharide (LPS) (Cat# L2630), adenosine triphosphate (ATP) (Cat# A7699), and phorbol-12-myristate-13-acetate (TPA) (Cat# P8139), were purchased from Sigma-Aldrich (St. Louis, MO, USA). Nigericin (Cat# tlrl-nig), Alum (Cat# tlrl-alk), Poly(dA:dT)/LyoVec (Cat# tlrl-patc), Lipofectamine 2000 (Cat# 11668019) were purchased from InvivoGene Biotech Co., Ltd. (San Diego, CA, USA).

X-α-Gal (Cat# XA250) was purchased from Gold Biotechnology (St. Louis, MO, USA). Aureobasidin A (Cat# 630499) was obtained from Clontech (Mountain View, CA, USA). Z-YVAD-FMK (Cat# 1012-100) was obtained from BioVision (Heinrichstr, Zürich). cOmplete, EDTA-free Protease Inhibitor Cocktail Tablets provided in EASYpacks (Cat# 04 693 132 001) was purchased form Roche (Indianapolis, IN, USA).

Anti-Human IL-1β (D3U3E) (Cat# 12703), Anti-Human Caspase-1 (D7F10) (Cat# 3866), Anti-Human/Mouse NLRP3 (D4D8T) (Cat# 15101), Anti-Human/Mouse/Rat Cyclin E1 (D7T3U) (Cat# 20808), Anti-Human/Mouse/Rat Skp1 (Cat# 2156), and Anti-Human/Mouse K63-linkage Specific Polyubiquitin (Cat# 5621) antibodies were purchased from Cell Signaling Technology (Beverly, MA, USA). Anti-Human/Mouse/Rat ASC(F-9) (Cat# sc-271054), Anti-Human/Mouse/Rat Cullin1 (D-5) (Cat# sc-17775), and Anti-Human/Mouse/Rat Cullin1 (H-213) (Cat# sc-11384) antibodies were purchased from Santa Cruz Biotechnology (Santa Cruz, CA, USA). Anti-Flag (Cat# F3165), Anti-HA (Cat# H6908), and Anti-GAPDH (Cat# G9295) antibodies were purchased from Sigma-Aldrich. Anti-Human/Mouse/Rat ROC1 (Cat# 14895-1-AP) antibody was obtained from Proteintech (Rosemont, IL, USA).

Anti-Human/Mouse NLRP3 (Cryo-2) (Cat# AG-20B-0014-C100) antibody was purchased from Adipogen (San Diego, CA, USA). Anti-Mouse IgG Dylight 649 (Cat# A23610), Anti-Mouse IgG Dylight 488 (Cat# A23210), Anti-Rabbit IgG Dylight 649 (Cat# A23620), and Anti-Rabbit IgG FITC (Cat# A22120) antibodies were purchased from Abbkine (California, USA). Human IL-1β ELISA Kit II (Cat# 557966) was purchased from BD Biosciences (Franklin Lakes, NJ, USA). Human IL-1β/IL-1F2 Quantikine ELISA Kit (Cat# DLB50) was purchased from R&D Systems (Minnesota, USA).

### Yeast two-hybrid screening

Mate & Plate Library-Universal Human (Normalized), Saccharomyces cerevisiae strain Y2HGold and Y187, control vectors pGBKT7, pGADT7, pGBKT7-p53, pGADT7-T and pGBKT7-lam, and some reagents were purchased from Clontech (Mountain View, CA, USA). All experimental procedures were following the Matchmaker® Gold Yeast Two-Hybrid System User Manual.

### Western blot analysis

For western blot analysis, cells were lysed in lyses buffer (50 mM Tris-HCl, pH7.4, 150 mM NaCl, 1% (vol/vol) Triton X-100, 5 mM EDTA, and 10% (vol/vol) glycerol). Protein concentration was determined by Bradford assay (Bio-Rad, Hercules, CA, USA). Cell lysates were separated by 10–12% SDS-PAGE and then transferred onto a nitrocellulose membrane (Millipore). The membranes were blocked in phosphate buffered saline with 0.1% Tween 20 (PBST) containing 5% nonfat dried milk before incubation with specific antibodies. Blots were detected with the Clarity Western ECL substrate (Bio-Rad, Hercules, CA, USA) and protein band were detected using a Luminescent image Analyzer (Fujifilm LAS-4000).

### Co-immunoprecipitation and immunoblot assays

Cells were washed with cold PBS and lysed in Nonidet P-40 lyses buffer (50 mM Tris-HCl, pH7.4, 150 mM NaCl, 1% (vol/vol) NP-40, 1 mM EDTA, and 5% (vol/vol) glycerol) containing protease inhibitors. A proportion of the lysates were saved for immunoblot analysis to detect the expression of target proteins. For immunoprecipitation, the rest of lysates were incubated with a control IgG or the indicated primary antibodies at 4°C for 12–16 h, and were further incubated with protein A/G-agarose (GE Healthcare, Milwaukee, WI, USA) for 3–4 h. The beads were washed for five times by 1ml washing buffer [50 mM Tris-HCl, pH7.4, 300 mM NaCl, 1% (vol/vol) NP-40, 1 mM EDTA, and 5% (vol/vol) glycerol] and reconstituted in 50 µl 2×SDS loading buffer before immunoblot analysis.

### Quantitative PCR

Total RNA was extracted with TRIzol reagent (Invitrogen, CA, USA), following the manufacturer’s instructions. Real-time quantitative RT-PCR was performed using the Roche LC480 and SYBR RT-PCR kits (DBI Bio-science, Ludwigshafen, Germany) in a reaction mixture of 10 μl SYBR Green PCR master mix, 1 μl DNA diluted template, 2 µl Real-time PCR primers and RNase-free water to complete the 20 μl volume. All Real-time PCR primers were designed in Nucleotide of National Center for Biotechnology Information (USA). All primers were as follows: GAPDH forward, 5’-AAGGCTGTGGGCAAGG-3’, GAPDH reverse, 5’-TGGAGGAGTGGGTGTCG-3’, IL-1β forward, 5’-CACGATGCACCTGTACGATCA-3’, IL-1β reverse, 5’-GTTGCTCCATATCCTGTCCCT-3’, Cullin1 forward, 5’-TTGCAAAGGGCCCTACGTT-3’, Cullin1 reverse 5’-CGTTGTTCCTCAAGCAGACG-3’, NLRP3 forward, 5’-AAGGGCCATGGACTATTTCC-3’, NLRP3 reverse, 5’-GACTCCACCCGATGACAGTT-3’, ASC forward, 5’-AACCCAAGCAAGATGCGGAAG-3, ASC reverse, 5-TTAGGGCCTGGAGGAGCAAG-3.

### Lentiviral production and infection

A pLKO.1-encoding shRNA vector for a scrambled (Sigma-Aldrich, St. Louis, MO, USA) or a specific-target molecule (Sigma-Aldrich, St. Louis, MO, USA) was transfected along with pMD2.G (an envelope plasmid) and psPAX2 (a packaging plasmid) into HEK293T cells. HEK293T culture supernatants were harvested at 36 and 48 h after transfection. Filtering the culture supernatants through a 0.45 µm filter to remove the cells. THP-1 cells and HEK293T cells were infected with collected culture supernatants plus 8 μg/ml Polybrene (Sigma-Aldrich, St. Louis, MO, USA) for 24 h. After 24 h, 1.5 μg/ml puromycin (InvivoGen, San Diego, CA, USA) was added into lentiviral-infected THP-1 cells for selection. 2 μg/ml puromycin (InvivoGen, San Diego, CA, USA) was added into lentiviral-infected HEK293T cells for selection. Knock down efficiency of each shRNA-targeted molecule was identified by RT-PCR and immunoblot analysis. The targeting sequences of shRNA for human Cullin1 are as follows: sh-Cullin1#1: 5’-GCACACAAGATGAATTAGCAA-3’, sh-Cullin1#2: 5’-GCCAGCATGATCTCCAAGTTA-3’. The targeting sequences of shRNA for human NLRP3 and ASC are as follows: sh-NLRP3: 5’-CAGGTTTGACTATCTGTTCT-3’, sh-ASC: 5’-GATGCGGAAGCTCTTCAGTTTCA-3’

### Reconstitution of the NLRP3 inflammasome in HEK293T cells

HEK293T cells at 70% confluence were seeded into 6cm plates overnight and then transfected with plasmids encoding NLRP3 inflammasome component proteins and pro-IL-1β, including 1 µg pcDNA3.1-NLRP3, 100 ng pcDNA3.1-ASC, 400 ng pcDNA3.1-procaspase-1 and 1 µg pcDNA3.1-pro-IL-1β. The cells were washed with culture medium 24 h after transfection and were further incubated for 6 h. Cell pellets were collected and lysed in cell lyses buffer for immunoblot analysis and Cell culture medium was collected for ELISA to detect IL-1β (BD Biosciences).

### Immunofluorescence microscopy

All Cells were washed three times with PBS, and cells were fixed with 4% paraformaldehyde for 15 min, washed three times with PBS, permeabilized with PBS containing 0.5% Triton X-100 for 5 min, washed three times with PBS, and finally blocked with PBS containing 5% bovine serum albumin (BSA) for 1 h at room temperature. The cells were then incubated with the primary antibody at 4°C, followed by incubation with FITC-conjugate donkey anti-mouse IgG (Abbkine) and Dylight 649-conjugate donkey anti-rabbit IgG (Abbkine) or FITC-conjugate donkey anti-rabbit IgG (Abbkine) and Dylight 649-conjugate donkey anti-mouse IgG (Abbkine) for 1 h, After cells washed three times with PBS, cells were incubated with DAPI (Vector Laboratories, Burlingame, CA) for 5 min in 37°C. Finally, the cells were analyzed using a confocal laser scanning microscope (FluoView FV 1000; Olympus, Tokyo, Japan).

### Cytokine measurements

The concentrations of human IL-1β in culture supernatants were measured by commercially available enzyme linked immunosorbent assay (ELISA) kits (BD Biosciences, San Jose, CA, USA). The concentrations of mouse IL-1β in peritoneal fluids or mouse IL-1β in the serum were measured by ELISA kits (R&D Systems, Minneapolis, USA). The concentrations of mouse IL-6 in peritoneal fluids were measured by ELISA kits (BioLegend, San Diego, USA).

### Measurement of activated Casp-1 and mature IL-1β

1 ml medium from each well was mixed with 0.5 ml methanol and 0.125 ml chloroform, vortexed, and centrifuged for 5 min at 13000 rpm in a microcentrifuge (Thermo scientific). The upper phase of each sample was removed and 0.5 ml methanol was added. Samples were centrifuged for 5 min at 13000 rpm. Next, the supernatants were removed and pellets were dried for 5 min at 50°C in the water bath. Then, 50 µl 2 x SDS loading buffer was added to each sample, followed by boiling for 10 min.

### *In vivo* ubiquitination assay

Cells were lysed in 100 µl SDS lyses buffer (50 mM Tris-HCl, pH7.4, 150 mM NaCl, 1% (vol/vol) NP-40, 1 mM EDTA, and 5% (vol/vol) glycerol, 1% SDS). After boiling for 5 min, lysates were diluted 10-fold with dilution buffer (50 mM Tris-HCl, pH7.4, 150 mM NaCl, 1% (vol/vol) NP-40, 1 mM EDTA, and 5% (vol/vol) glycerol) containing protease inhibitors. A proportion of the lysates (100 µl) were saved for immunoblot analysis to detect the expression of target proteins, the rest of the lysates were immunoprecipitated with specific antibodies. Rest of experiment procedures were following the co-immunoprecipitation and immunoblot assays.

### Statistics

All experiments were repeated at least three times with similar results. All results were expressed as the mean ± the standard deviation (SD). Statistical analysis was carried out using the t-test for two groups and one-way ANOVA for multiple groups (GraphPad Prism5). The date was considered statistically significant when P ≤0.05 (*), P ≤0.01 (**), P ≤0.001 (***).

## Acknowledgments

This work was supported by National Natural Science Foundation of China (81730061, 31230005, 81471942, 31200134, and 31270206), and the National Health and Family Planning Commission of China (National Mega Project on Major Infectious Disease Prevention) (2017ZX10103005 and 2017ZX10202201).

We thank Dr. Jiahuai Han of State Key Laboratory of Cellular Stress Biology, School of Life Sciences, Xiamen University, Fujian, China, for kindly providing the cDNA of genes Cullin1 and NLRP3.

## Author Contributions

P.W., Q.Z., Y.L., K.W., Y.L., and J.W. conceived the project and designed the experiments; P.W. performed experiments with help from Q.Z., W.L., Y.J., T.W., W.W., P.P., G.Y., Q.X., S.H., Q.Y., and W.Z.; W.L., and F.L., contributed to the reagents; P.W., Q.Z., W.L., W.W., K.W., F.L., Y.L., and J.W. performed data analyses; P.W., and J.W. wrote the manuscript; P.W., K.W., Y.L., and J.W. edited the manuscript.

## Conflict of Interest

The authors disclose no conflicts or competing financial interests.

## Expanded View Figure Legends

**Figure EV1 – Two-hybrid library screening using yeast mating.**

The entire Matchmaker screening process consists of the following steps: Step 1, Clone the targeted gene into pGBKT7 Vector; Step 2, Test bait for autoactivation and toxicity; Step3, Screen Mate & Plate library; Step 4, confirm and interpret results.

**Figure EV2 – Analyses of the efficiency of short hairpin RNAs (shRNA) in HEK293T cells.**

**A** HEK293T cells were transfected with pcDNA3.1(+)-3XFlag-Cullin1 along with control, plasmid pSilencer2.1-U6-shCullin1#1 and shCullin1#2. The levels of Flag-Cullin1 protein were detected Western blot analyses.

**B** HEK293T cells were transfected with pcDNA3.1(+)-3XFlag-Cullin1 along with control, plasmid pSilencer2.1-U6-shCullin1#1 and shCullin1#2. The levels of ROC1 protein were detected Western blot analyses.

**C** HEK293T cells were transfected with pcDNA3.1(+)-3XFlag-Cullin1 along with control, plasmid pSilencer2.1-U6-shCullin1#1 and shCullin1#2. The levels of SKP1 protein were detected Western blot analyses.

The method of verifying knockdown its gene of pSilencer2.1-U6-shROC1 or pSilencer2.1-U6-shSKP1 was similar to that above.

**Figure EV3 – The effect of IAV H3N2 on the activation of IL-1 and the role of Cullin1 in regulating the components of NLRP3 inflammasome.**

**A** PMA-differentiated THP-1 macrophage cells were treated by H3N2 for 24 h and different concentration of Z-YVAD-FMK (inhibitor of Caspase-1) for 6 h. Supernatants were analyzed by ELISA for IL-1β secretion.

**B** PMA-differentiated THP-1 macrophage cells were treated by H3N2 for 24 h and different concentration of Z-YVAD-FMK (inhibitor of Caspase-1) for 6 h. The level of IL-1β mRNA was determined by qRT-PCR.

**C** THP-1 cells were transduced with lentiviruses stably expressing shRNA(shNC) and shRNA against NLRP3 (sh-NLRP3) and ASC (sh-ASC) and selected with puromycin for 2 weeks. The levels of NLRP3 mRNA were determined by qRT-PCR.

**D** THP-1 cells were transduced with lentiviruses stably expressing shRNA(shNC) and shRNA against NLRP3 (sh-NLRP3) and ASC (sh-ASC) and selected with puromycin for 2 weeks. The levels of ASC mRNA were determined by qRT-PCR.

**E** TPA-differentiated THP-1 cells with shRNA mediated knockdown of gene expression were infected by influenza A virus H3N2, the supernatants were analyzed by ELISA for IL-1β secretion.

**F** The level of IL-1β mRNA was determined by qRT-PCR in THP-1 cells and PMA-differentiated THP-1 cells.

**G** THP-1 cells were transducted with lentiviruses stably expressing Cullin1 and selected with puromycin for 2 weeks. The level of Cullin1 and GAPDH proteins were detected by RT PCR and Western blot analyses.

**H** THP-1 cells were transducted with lentiviruses stably expressing shRNA(sh-NC) and shRNA against Cullin1 (sh-Cullin1#1 and sh-Cullin1#2) and selected with puromycin for 2 weeks. The level of Cullin1 and GAPDH mRNA was determined by qRT-PCR(upper panel) and the level of Cullin1 and GAPDH proteins were detected Western blot analyses (lower panel).

**I** Proteins level of Cullin1, Cyclin E1 and GAPDH were examined by Western blot analysis in THP-1 cells stably expressing sh-NC and shCullin#2.

**Figure EV4 – Identification of interaction between ASC and the PYD of NLRP3.**

HEK293T cells were transfected with HA-ASC plasmid in combination with plasmids encoding Flag-PYD, Flag-NBD and Flag-LRR. The cell lysates were immunoprecipitated with control IgG or an anti-Flag antibody and then immunoblotted with the indicated antibodies.

**Figure EV5 – Cullin1 can’t promote degradation of NLRP3.**

HEK293T cells were transfected with pCAGGS-HA-NLRP3 along with control and three different quantities of pcDNA3.1(+)-3XFlag-Cullin. Lysates were immunoblotted with the indicated antibodies.

## References

1. Meylan E, Tschopp J, Karin M (2006) Intracellular pattern recognition receptors in the host response. Nature 442: 39–44.

2. Takeuchi O, Akira S (2010) Pattern recognition receptors and inflammation. Cell 140: 805– 820.

3. Ting JP, Duncan JA, Lei Y (2010) How the noninflammasome NLRs function in the innate immune system. Science 327: 286–290.

4. Fink SL, Bergsbaken T, Cookson BT (2008) Anthrax lethal toxin and Salmonella elicit the common cell death pathway of caspase-1-dependent pyroptosis via distinct mechanisms. Proc Natl Acad Sci USA 105: 4312–4317.

5. Martinon F, Burns K, Tschopp J (2002) The inflammasome: a molecular platform triggering activation of inflammatory caspases and processing of proIL-beta. Mol Cell 10: 417–426.

6. Martinon F, Mayor A, Tschopp J (2009) The inflammasomes: guardians of the body. Ann Rev Immunol 27: 229–265.

7. Schroder K, Tschopp J (2010) The inflammasomes. Cell 140: 821–832.

8. Sutterwala FS, Ogura Y, Flavell RA (2007) The inflammasome in pathogen recognition and inflammation. J Leukoc Biol 82: 259–264.

9. Srinivasula SM, Poyet JL, Razmara M, Datta P, Zhang Z, Alnemri ES (2002) The PYRIN-CARD protein ASC is an activating adaptor for caspase-1. J Biol Chem 277: 21119–21122.

10. Vajjhala PR, Mirams RE, Hill JM (2012) Multiple binding sites on the pyrin domain of ASC protein allow self-association and interaction with NLRP3 protein. J Biol Chem 287: 41732–41743.

11. Black RA, Kronheim SR, Merriam JE, March CJ, Hopp TP (1989) A pre-aspartate-specific protease from human leukocytes that cleaves pro-interleukin-1 beta. J Biol Chem 264: 5323–5326.

12. Cerretti DP, Kozlosky CJ, Mosley B, Nelson N, Van Ness K, Greenstreet TA, March CJ, Kronheim SR, Druck T, Cannizzaro LA, et al (1992) Molecular cloning of the interleukin-1 beta converting enzyme. Science 256: 97–100.

13. Kostura MJ, Tocci MJ, Limjuco G, Chin J, Cameron P, Hillman AG, Chartrain NA, Schmidt JA (1989) Identification of a monocyte specific pre-interleukin 1 beta convertase activity. Proc Natl Acad Sci USA 86: 5227–5231.

14. Thornberry NA, Bull HG, Calaycay JR, Chapman KT, Howard AD, Kostura MJ, Miller DK, Molineaux SM, Weidner JR, Aunins J, et al (1992) A novel heterodimeric cysteine protease is required for interleukin-1 beta processing in monocytes. Nature 356: 768–774.

15. Winkler S, Rosen-Wolff A (2015) Caspase-1: an integral regulator of innate immunity. Semin Immunopathol 37: 419–427.

16. Fernandes-Alnemri T, Wu J, Yu JW, Datta P, Miller B, Jankowski W, Rosenberg S, Zhang J, Alnemri ES (2007) The pyroptosome: a supramolecular assembly of ASC dimers mediating inflammatory cell death via caspase-1 activation. Cell Death Differ 14: 1590–1604.

17. Latz E, Xiao TS, Stutz A (2013) Activation and regulation of the inflammasomes. Nat Rev Immunol 13: 397–411.

18. Petrilli V, Papin S, Tschopp J (2005) The inflammasome. Curr Biol 15: R581.

19. Proell M, Gerlic M, Mace PD, Reed JC, Riedl SJ (2013) The CARD plays a critical role in ASC foci formation and inflammasome signalling. Biochem J 449: 613–621.

20. Masters SL, Dunne A, Subramanian SL, Hull RL, Tannahill GM, Sharp FA, Becker C, Franchi L, Yoshihara E, Chen Z, et al (2010) Activation of the NLRP3 inflammasome by islet amyloid polypeptide provides a mechanism for enhanced IL-1beta in type 2 diabetes. Nat Immunol 11: 897–904.

21. Strowig T, Henao-Mejia J, Elinav E, Flavell R (2012) Inflammasomes in health and disease. Nature 481: 278–286.

22. Sutterwala FS, Ogura Y, Szczepanik M, Lara-Tejero M, Lichtenberger GS, Grant EP, Bertin J, Coyle AJ, Galán JE, Askenase PW, et al (2006) Critical role for NALP3/CIAS1/Cryopyrin in innate and adaptive immunity through its regulation of caspase-1. Immunity 24: 317–327.

23. Walsh JG, Reinke SN, Mamik MK, McKenzie BA, Maingat F, Branton WG, Broadhurst DI, Power C (2014) Rapid inflammasome activation in microglia contributes to brain disease in HIV/AIDS. Retrovirology 11: 35.

24. Wen H, Ting JP, O’Neill LA (2012) A role for the NLRP3 inflammasome in metabolic diseases-did Warburg miss inflammation? Nat Immunol 13: 352–357.

25. Willingham SB, Bergstralh DT, O’Connor W, Morrison AC, Taxman DJ, Duncan JA, Barnoy S, Venkatesan MM, Flavell RA, Deshmukh M, et al (2007) Microbial pathogen-induced necrotic cell death mediated by the inflammasome components CIAS1/cryopyrin/NLRP3 and ASC. Cell Host Microbe 2: 147–159.

26. Johnston JB, Barrett JW, Nazarian SH, Goodwin M, Ricciuto D, Wang G, McFadden G (2005) A poxvirus-encoded pyrin domain protein interacts with ASC-1 to inhibit host inflammatory and apoptotic responses to infection. Immunity 23: 587598.

27. Yan Y, Jiang W, Liu L, Wang X, Ding C, Tian Z, Zhou R (2015) Dopamine controls systemic inflammation through inhibition of NLRP3 inflammasome. Cell 160: 62–73.

28. Yang CS, Kim JJ, Kim TS, Lee PY, Kim SY, Lee HM, Shin DM, Nguyen LT, Lee MS, Jin HS, et al (2015) Small heterodimer partner interacts with NLRP3 and negatively regulates activation of the NLRP3 inflammasome. Nat Commun 6: 6115.

29. Chondrogianni N, Gonos ES. (2012) Structure and function of the ubiquitin-proteasome system: modulation of components. Prog Mol Biol Transl Sci 109: 41–74.

30. Finley D (2009) Recognition and processing of ubiquitin-protein conjugates by the proteasome. Ann Rev Biochem 78: 477–513.

31. Glickman MH, Ciechanover A (2002) The ubiquitin-proteasome proteolytic pathway: destruction for the sake of construction. Physiol Rev 82: 373–428.

32. Sun Y (2006) E3 ubiquitin ligases as cancer targets and biomarkers. Neoplasia 8: 645–654.

33. Deshaies RJ, Joazeiro CA (2009) RING domain E3 ubiquitin ligases. Ann Rev Biochem 78: 399–434.

34. Burrows AC, Prokop J, Summers MK (2012) Skp1-Cul1-F-box ubiquitin ligase (SCF(betaTrCP))-mediated destruction of the ubiquitin-specific protease USP37 during G2-phase promotes mitotic entry. J Biol Chem 287: 39021–39029.

35. Hershko A, Ciechanover A (1998) The ubiquitin system. Ann Rev Biochem 67: 425–479.

36. Zheng N, Schulman BA, Song L, Miller JJ, Jeffrey PD, Wang P, Chu C, Koepp DM, Elledge SJ, Pagano M, et al (2002) Structure of the Cul1-Rbx1-Skp1-F boxSkp2 SCF ubiquitin ligase complex. Nature 416: 703–709.

37. Bai C, Sen P, Hofmann K, Ma L, Goebl M, Harper JW, Elledge SJ (1996) SKP1 connects cell cycle regulators to the ubiquitin proteolysis machinery through a novel motif, the F-box. Cell 86: 263–274.

38. Skowyra D, Craig KL, Tyers M, Elledge SJ, Harper JW (1997) F-box proteins are receptors that recruit phosphorylated substrates to the SCF ubiquitin-ligase complex. Cell 91: 209– 219.

39. Bosu DR, Kipreos ET (2008) Cullin-RING ubiquitin ligases: global regulation and activation cycles. Cell Division 3: 7.

40. Nakayama KI, Nakayama K (2006) Ubiquitin ligases: cell-cycle control and cancer. Nat Rev Cancer 6: 369–381.

41. Petroski MD, Deshaies RJ (2005) Function and regulation of cullin-RING ubiquitin ligases. Nat Rev Mol Cell Biol 6: 9–20.

42. Sarikas A, Hartmann T, Pan ZQ (2011) The cullin protein family. Genome Biol 12(4): 220.

43. Bai J, Zhou Y, Chen G, Zeng J, Ding J, Tan Y, Zhou J, Li G (2011) Overexpression of Cullin1 is associated with poor prognosis of patients with gastric cancer. Human Pathol 42: 375–383.

44. Bai J, Yong HM, Chen FF, Mei PJ, Liu H, Li C, Pan ZQ, Wu YP, Zheng JN (2013) Cullin1 is a novel marker of poor prognosis and a potential therapeutic target in human breast cancer. Ann Oncol 24: 2016–2022.

45. Chen G, Cheng Y, Martinka M, Li G (2010) Cul1 expression is increased in early stages of human melanoma. Pigment Cell Melanoma Res 23: 572–574.

46. Chen G, Li G (2010) Increased Cul1 expression promotes melanoma cell proliferation through regulating p27 expression. International J Oncol 37: 1339–1344.

47. Dealy MJ, Nguyen KV, Lo J, Gstaiger M, Krek W, Elson D, Arbeit J, Kipreos ET, Johnson RS (1999) Loss of Cul1 results in early embryonic lethality and dysregulation of cyclin E. Nat Genet 23: 245–248.

48. Lu B, Nakamura T, Inouye K, Li J, Tang Y, Lundbäck P, Valdes-Ferrer SI, Olofsson PS, Kalb T, Roth J, et al (2012) Novel role of PKR in inflammasome activation and HMGB1 release. Nature 488: 670–674.

49. Moriyama M, Chen IY, Kawaguchi A, Koshiba T, Nagata K, Takeyama H, Hasegawa H, Ichinohe T (2016) The RNA- and TRIM25-binding domains of influenza virus NS1 protein are essential for suppression of NLRP3 inflammasome-mediated interleukin-1beta secretion. J Virol 90: 4105–4114.

50. Shi H, Wang Y, Li X, Zhan X, Tang M, Fina M, Su L, Pratt D, Bu CH, Hildebrand S, et al (2016) NLRP3 activation and mitosis are mutually exclusive events coordinated by NEK7, a new inflammasome component. Nat Immunol 17(3): 250–258.

51. Wang W, Li G, Wu D, Luo Z, Pan P, Tian M, Wang Y, Xiao F, Li AX, Wu K, et al (2018) Zika virus infection induces host inflammatory responses by facilitating NLRP3 inflammasome assembly and interleukin-1β secretion. Nat Commun 9: 106.

52. Hao J, Song X, Wang J, Guo C, Li Y, Li B, Zhang Y, Yin Y (2015) Cancer-testis antigen MAGE-C2 binds Rbx1 and inhibits ubiquitin ligase-mediated turnover of cyclin E. Oncotarget 6: 42028–42039.

53. Welcker M, Clurman BE (2008) FBW7 ubiquitin ligase: a tumour suppressor at the crossroads of cell division, growth and differentiation. Nat Rev Cancer 8: 83–93.

54. Elliott EI, Sutterwala FS (2015) Initiation and perpetuation of NLRP3 inflammasome activation and assembly. Immunol Rev 265: 35–52.

55. Tschopp J, Schroder K (2010). NLRP3 inflammasome activation: The convergence of multiple signalling pathways on ROS production? Nat Rev Immunol 10: 210–215.

56. Rathinam VA, Vanaja SK, Fitzgerald KA (2012) Regulation of inflammasome signaling. Nat Immunol 13: 333–342.

57. Song H, Liu B, Huai W, Yu Z, Wang W, Zhao J, Han L, Jiang G, Zhang L, Gao C, et al (2016) The E3 ubiquitin ligase TRIM31 attenuates NLRP3 inflammasome activation by promoting proteasomal degradation of NLRP3. Nat Commun 7: 13727.

58. Juliana C, Fernandes-Alnemri T, Kang S, Farias A, Qin F, Alnemri ES (2012) Non-transcriptional priming and deubiquitination regulate NLRP3 inflammasome activation. J Biol Chem 287: 36617–36622.

59. Py BF, Kim MS, Vakifahmetoglu-Norberg H, Yuan J (2013) Deubiquitination of NLRP3 by BRCC3 critically regulates inflammasome activity. Mol Cell 49: 331–338.

60. Radivojac P, Vacic V, Haynes C, Cocklin RR, Mohan A, Heyen JW, Goebl MG, Iakoucheva LM (2010) Identification, analysis, and prediction of protein ubiquitination sites. Proteins 78: 365–380.

61. Guarda G, Braun M, Staehli F, Tardivel A, Mattmann C, Förster I, Farlik M, Decker T, Du Pasquier RA, Romero P, et al (2011) Type I interferon inhibits interleukin-1 production and inflammasome activation. Immunity 34(2): 213–223.

62. Latz E (2010) The inflammasomes: mechanisms of activation and function. Curr Opin Immunol 22: 28–33.

63. Leemans JC, Cassel SL, Sutterwala FS (2011) Sensing damage by the NLRP3 inflammasome. Immunol Rev 243: 152–162.

64. Prochnicki T, Mangan MS, Latz E (2016) Recent insights into the molecular mechanisms of the NLRP3 inflammasome activation. F1000Res 5: pii: F1000 Faculty Rev-1469.

65. Furukawa M, Zhang Y, McCarville J, Ohta T, Xiong Y (2000) The CUL1 C-terminal sequence and ROC1 are required for efficient nuclear accumulation, NEDD8 modification, and ubiquitin ligase activity of CUL1. Mol Cell Biol 20: 8185–8197.

66. Park S, Juliana C, Hong S, Datta P, Hwang I, Fernandes-Alnemri T, Yu JW, Alnemri ES (2013) The mitochondrial antiviral protein MAVS associates with NLRP3 and regulates its inflammasome activity. J Immunol 191, 4358–4366.

67. Subramanian N, Natarajan K, Clatworthy MR, Wang Z, Germain RN (2013) The adaptor MAVS promotes NLRP3 mitochondrial localization and inflammasome activation. Cell 153: 348–361.

68. Han S, Lear TB, Jerome JA, Rajbhandari S, Snavely CA, Gulick DL, Gibson KF, Zou C, Chen BB, Mallampalli RK (2015) Lipopolysaccharide primes the NALP3 inflammasome by inhibiting its ubiquitination and degradation mediated by the SCFFBXL2 E3 ligase. J Biol Chem 290: 18124–18133.

